# Circadian rhythms in the gut microbiota shape sex differences in host gene expression and metabolism

**DOI:** 10.1101/2022.09.15.508006

**Authors:** Sarah K. Munyoki, Julie P. Goff, Antonija Kolobaric, Armari Long, Steven J. Mullett, Jennifer K. Burns, Aaron K. Jenkins, Lauren DePoy, Stacy G. Wendell, Colleen A. McClung, Kathleen E. Morrison, Eldin Jašarević

## Abstract

Circadian rhythms dynamically regulate sex differences in metabolism and immunity, and circadian disruption increases the risk of metabolic disorders. We investigated the role of sex-specific microbial circadian rhythms in host metabolism using germ-free and conventionalized female and male mice, dietary manipulations, coupled with a systems biology approach. Sex differences in circadian rhythms of genes involved in immunity and metabolism are dependent on oscillations in the microbiota, microbial metabolic functions, and microbial metabolites. Further, dietary factors modify the magnitude of sex differences in host-microbe circadian dynamics. We show that consuming an obesogenic high-fat, low-fiber diet produced sex-specific changes in circadian rhythms in microbiota, metabolites, and host gene expression, which were linked to sex differences in the severity of metabolic dysfunction. These results reveal that microbial circadian rhythms contribute to sex differences in metabolism, emphasizing the need to consider sex as a biological variable in research on microbial contributions to metabolic dysfunction.

**HIGHLIGHTS:**

- Microbial circadian rhythms differ by sex.
- Sex-specific rhythms in host transcriptional networks are microbiome-dependent.
- Diet-induced obesity entrains new sex-specific rhythms in microbiome and host genes.
- Timing of data collection influences magnitude of sex differences.

## INTRODUCTION

Sex differences in physiologic, metabolic, immune, and behavioral processes are well-documented across all vertebrate species, including humans^1–7^. The maintenance of these sex differences depends on a complex interplay between genetic, environmental, and developmental factors^1, 2^. One consequence of these biological sex differences is that most adult-onset diseases exhibit a sex specific bias in prevalence, age of onset, severity, and treatment outcome^6^. For instance, autoimmunity, obesity, metabolic syndrome, and type 2 diabetes exhibit significant sex differences in disease management and treatment response to dietary and pharmacological interventions. Despite the clear links between sex-specific physiology and disease trajectories, the historical underrepresentation of women in these studies has created a barrier to uncovering underlying mechanisms that may provide novel insights into disease prevention, intervention, and treatment.

Microbial communities within the intestinal tract are key players in balancing metabolic homeostasis and dysfunction^8, 9^. At birth, microbial composition is similar between female and male mice, but around puberty, these communities begin to diverge until they reach steady-state sex differences in adulthood^10^. Although the adult intestinal microbiota is typically stable, recent studies suggest that the relative abundance of microbiota varies significantly throughout the day, coordinating with the circadian clock and feeding patterns to generate diurnal rhythms in energy homeostasis, lipid metabolism, innate immune function, and energy substrate provision to distal tissues^11–22^. Moreover, individual variability in environmental and lifestyle exposures, including jet lag, shift work, antibiotic exposure, altered feeding times and o consumption of highly processed diets, are associated with disruption to diurnal rhythms in gut microbiota ^20, 25–29^. In turn, microbial circadian misalignment is reported to increase the risk for obesity, poor glycemic control, metabolic dysfunction, inflammation, and overall heightened susceptibility to disease ^30–34^.

Peripheral circadian rhythms also influence the composition of microbial communities in a sex-specific manner. The total bacterial load and microbial composition significantly change throughout the day, and these oscillations are more pronounced in female mice than male mice ^23^. Deletion of *Bmal1*, the principal driver of the mammalian molecular clock, abolishes sex differences in the microbiota composition^23^. Sex differences in hepatic gene expression, metabolism, and reproductive development are disrupted in germ-free mice, suggesting that microbial-derived signals are necessary for sex-specific development and phenotype ^24^. Although circadian disruption is often associated with sex-specific health outcomes, most studies on circadian rhythms, the microbiome, and health outcomes often exclude female mice from studies altogether^35^.

This study aimed to identify sex-specific circadian rhythms in the microbial, metabolic, and transcriptional capacity under *ad libitium* feeding conditions. We used germ-free, conventionalized and murine-pathogen free mice, and dietary manipulations to examine the central hypothesis that circadian rhythms in diet-host-microbe interactions differ between female and male mice. Despite well-established observation that females are resistant to diet-induced obesity^3636, 37^, female mice are currently excluded in studies on circadian rhythms, diet-induced obesity, and the microbiome. To address this fundamental gap, we investigated whether circadian disruption caused by consumption of a high-fat low-fiber diet affects the microbiome, microbial metabolites, and host transcriptome in a sex-specific manner. To examine these hypotheses, we employed an integrated multi-Omics approach to identify sex differences in diurnal dynamics of the intestinal microbiota, microbial derived metabolite production, systemic availability of microbial metabolites, and host transcriptome.

## RESULTS

### Dietary factors link sex-specific circadian rhythms in gut microbiota composition to severity of metabolic dysfunction

Consumption of a high-fat low-fiber (Hf-Lf) diet produces hallmark phenotypes of diet-induced obesity and metabolic syndrome, including increased rapid weight gain, excess accumulation of fat mass, hyperglycemia, hyperinsulinemia, metabolic endotoxemia, and low-grade chronic inflammation^38–41^. A more severe diet-induced metabolic phenotype is observed in male mice, while protection or resistance to diet-induced obesity is observed in female mice^42, 43^. Hf-Lf feeding alters gut microbiota circadian rhythms in male mice, but the role of Hf-Lf-diet induced obesity on female specific microbial rhythms is unknown^26–28^. To investigate possible sex differences, we randomly assigned pubertal female and male mice to either an Hf-Lf diet (20% protein, 20% carbohydrate, 60% fat) or a chow diet (23.2% protein, 55.2% carbohydrate, 21.6% fat)^44, 45^ (**Figure 1A**). It is important to note that the lack of complex carbohydrates and soluble fibers in commercially available refined high-fat diet formulations (e.g., Research Diets OpenSource Diets) is proposed to be a contributing factor to excessive weight gain, obesity, and diabetes in mouse models of diet induced metabolic syndrome and obesity. Hence, the dietary designation Hf-Lf serves to highlight absence of microbiota accessible carbohydrates in these dietary formulations^45–50^.

**Figure 1.**
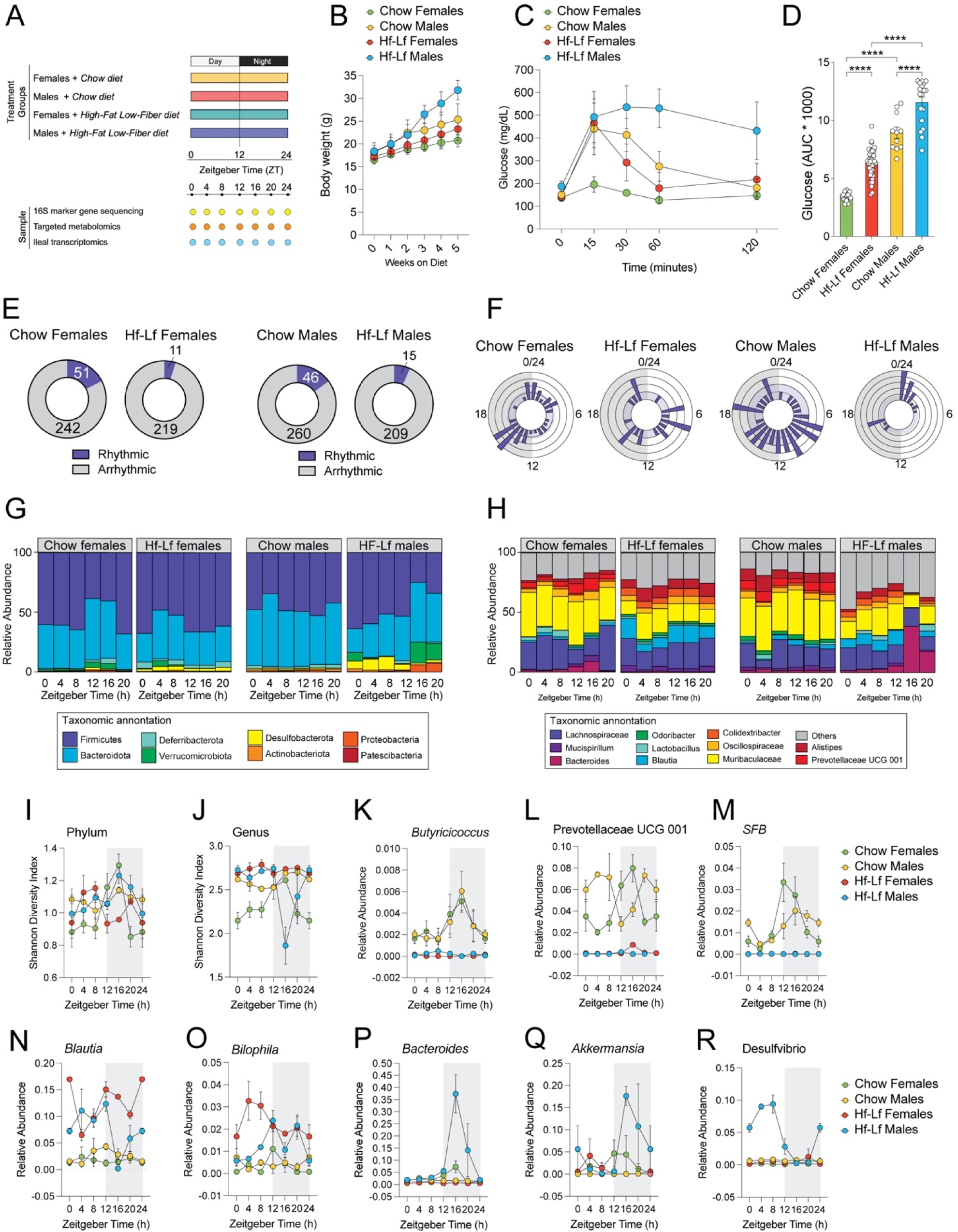
Diet modifies sex differences in diurnal rhythmicity of the intestinal microbiota. A) Schematic representation of study design and sample collections. B) Sex-specific effects of Chow and Hf-Lf diet on body weight gain (time*sex interaction, F_15, 419_ = 38.75, *<* 0.0001). Hf-Lf females weigh more than Chow females (*t*_84_ = 5.37, *p* = 0.0016) and Hf-Lf males weigh more than Chow males (*t*_84_ = 4.041, *p* = 0.027). Chow males weigh more than Hf-Lf females (*t*_84_ = 4.973, *p* = 0.004), showing that Hf-Lf female mice weigh less than Chow males even after 6wks on Hf-Lf feeding. C) Sex-specific effects of Chow and Hf-Lf diet in glucose tolerance (time*sex interaction, F_12,316_ = 33.82, *p <* 0.0001). Hf-Lf females showed slower glucose clearance than Chow females (*t*_79_ = 10.15, *p* < 0.0001) and Hf-Lf males showed slower glucose clearance than Chow males (*t*_84_ = 4.973, *p* = 0.004). Chow males showed similar glucose clearance as Hf-Lf females (*t*_79_ = 2.803, *p* = 0.20). C) Sex-specific effects of Chow and Hf-Lf diet on circulating glucose levels (main effect of treatment, F_3,79_ = 107.7, *p <* 0.0001). Hf-Lf females had higher glucose levels than Chow females (*t*_79_ = 9.73, *p* < 0.0001) and Hf-Lf males had higher glucose levels than Chow males (*t*_79_ = 7.005, *p* < 0.0001). Chow males had higher glucose levels than Hf-Lf females (*t*_79_ = 7.835, *p* < 0.0001). E) Sex differences in non-rhythmic and rhythmic microbiota detected in the cecum using cosinor analysis (Cosinor *p* < 0.06). F) Rose plot showing sex differences in acrophase distribution of microbiota in the cecum of adult females (left) and males (right). G) Relative abundance of top eight phyla across a 24-hr period in Chow and Hf-Lf mice. H) Relative abundance of top twelve genera across a 24-hr period in Chow and Hf-Lf mice. I) Phylum-level alpha diversity differed by time-of-day in males and females. J) Genus-level alpha diversity differed by time-of-day in males and females. K) Diurnal variation in *Butyricicoccus* relative abundance detected in Chow mice but lost in Hf-Lf mice. L) Diurnal variation in *Prevotellaceae UCG 001* relative abundance detected in Chow mice but lost in Hf-Lf mice. M) Diurnal variation in Segmented Filamentous Bacteria relative abundance detected in Chow mice but lost in Hf-Lf mice. N) Emergence of diurnal variation in *Blautia* relative abundance detected in Hf-Lf females relative to Hf-Lf males and Chow mice. O) Emergence of diurnal variation in *Bilophila* relative abundance detected in Hf-Lf females relative to Hf Lf males and Chow mice. P) Emergence of diurnal variation in *Bacteroides* relative abundance detected in Hf-Lf males relative to Hf Lf females and Chow mice. Q) Emergence of diurnal variation in *Akkermansia* relative abundance detected in Hf-Lf males relative to Hf-Lf females and Chow mice. R) Emergence of diurnal variation in *Desulfovibrio* relative abundance detected in Hf-Lf males relative to Hf-Lf females and Chow mice. Three murine-pathogen free C57Bl/6NTac females and males were used for each condition, for a total of 36 females and 36 males. Acrophases were calculated by cosinor analysis (period = 24 h). Threshold for the cosinor test was set to *p < 0.05*. Full cosinor analysis in **TableS 1**. RM or One-way ANOVA, followed by Tukey correction for multiple testing. Data is represented as mean ± SEM. \**p*lJ<lJ0.05, ** *p <*lJ0.01, *** *p <*lJ0.001.

To confirm diet-induced metabolic syndrome, mice were given *ad libitum* access to their respective diets throughout the experiment, and weekly body weights were recorded. Glycemic control was evaluated in adulthood using an intraperitoneal glucose tolerance test (GTT). As expected, Hf-Lf mice showed sex-specific weight gain and impaired glucose clearance compared to Chow mice (**Figure 1B-D**). Consistent with a resistance to Hf-Lf-induced obesity that is specific to females, Hf-Lf female mice weighed less and showed a similar glucose clearance response compared to Chow males, even after six weeks of Hf-Lf feeding (**Figure 1B-D**).

We next examined whether the sex-specific weight gain and glucose intolerance caused by Hf-Lf diet affected diurnal variations in the gut microbiota differently in female and male mice. We collected cecal luminal contents from female and male mice that had been assigned to either a Chow or Hf-Lf diet and collected samples every 4 hours for 24 hours. We then analyzed the microbiota in these samples using 16S rRNA marker gene sequencing and applied cosinor analysis to identify sex-specific rhythms in microbiota^51^. Our analysis revealed that 54 out of 293 (∼17%) taxa showed oscillations in Chow females, and 46 out of 306 (∼15%) taxa did so in Chow males (all *p*s < 0.05 by Cosinor test). Only 14 taxa exhibited similar patterns in both Chow female and male mice (**Figure 1E, TableS1**). In Hf-Lf mice, the number of rhythmic taxa decreased to ∼4% and ∼6.5% in females and males, respectively. Analysis of microbial acrophases, defined as the peak relative abundance of each taxon, revealed that the distribution of acrophases was localized to two time periods in Chow females: Zeitgeber Time (ZT)12 to ZT18, and ZT0 to ZT4 (where ZT0 is lights on and ZT12 is lights off). In contrast, acrophase distribution in Chow males extended across multiple time points (**Figure 1E**). Hf-Lf feeding disrupted the sex-specific distribution of acrophases present in Chow mice (**Figure 1F**), indicating that diet may play a role in influencing the phase distribution of oscillating microbiota in a sex-specific manner.

We then sought to identify how diet influences oscillations of specific taxa at the phylum and genus level in female and male mice. At the phylum level, the relative abundance of Bacteroidota, Firmicutes, Verrucomicrobiota, and Deferribacterota showed significant variation at ZT12 and ZT16 in Chow-fed female and male mice (**Figure 1G**) ^23^. Hf-Lf reduced relative abundance of Firmicutes in both sexes, while Hf-Lf males showed a sex-specific expansion of Desulfobacterota during the behaviorally inactive phase (ZT0 and ZT12). This expansion was replaced by a bloom in the relative abundance of Bacteroidota, Proteobacteria, and Verrucomicrobiota during the behaviorally active phase (ZT16 and ZT20). These sex and diet-specific effects were similarly detected at the genus level and influenced community alpha diversity, as measured by the Shannon Diversity Index (**Figure 1H-J**).

Analysis at the genus level revealed additional sex-specific effects of diet on diurnal variation of microbiota composition (**Figure 1G; TableS1**). *Butyricicoccus*, a well-characterized butyrate producer, showed similar rhythmicity in Chow-fed males and females, with peak abundance around ZT16 (**Figure 1K**). *Segmented Filamentous Bacteria* showed sex-specific rhythmicity, with peak abundance occurring in females prior to males (ZT12 vs. ZT16, respectively) (**Figure 1M**). The relative abundance of *Mucispirillum* showed sex-specific rhythmicity in Chow-fed mice, with peak abundance in males at ZT20. *Prevotellaceae* UCG 001 showed sex-specific rhythmicity in Chow-fed mice, whereby relative abundance was in anti-phase in Chow-fed males and females (**Figure 1L**). *Alistipes* showed a Chow-fed male-specific increase during the behaviorally inactive phase but not in Chow-fed females. *Lactobacillus* showed peak abundance during the light phase, with Chow-fed females showing a higher overall abundance of this taxon. The relative abundance of *Muribaculaceae* (formerly S24-7) was stable across the time of day in Chow-fed males and females (all Cosnior p’s > 0.05). Rhythmicity in *Butyricicoccus*, *Prevotellaceae* UCG 001, and *Segmented Filamentous Bacteria* was abolished in Hf-Lf female and male mice, suggesting that feeding rhythms and dietary factors within the Chow diet synergize to entrain rhythms of these taxa (Cosinor *ps* > 0.05) (**Figure 1K-M**).

Loss of oscillations in the relative abundance of microbiota occurred concomitantly with a sex-specific gain of rhythm in Hf-Lf mice. Hf-Lf females gained rhythmicity in the relative abundance of *Blautia* and *Bilophila* compared with Hf-Lf males and Chow mice (Cosinor *p*s < 0.05) (**Figure 1N,O**). Hf-Lf males showed a gain of rhythmicity in the relative abundance of *Bacteroides*, *Akkermansia*, and *Desulfovibrio* compared to the other mice (Cosinor *p*s < 0.05) (**Figure 1P,Q,R**). Our results suggest that nutrients derived from diets that vary in carbohydrates, fat, and soluble fiber may exert sex-specific effects on diurnal variations in microbial structure, diversity, and composition within the cecal luminal contents of mice.

### Diet drives sex-specific circadian rhythms in predicted microbial functional pathways

Sex differences in diurnal variation of microbiota composition suggest that core metabolic functions of the microbiome may also vary depending on the time of day. We conducted a gut metabolic modules (GMM) analysis to determine whether sex-specific oscillations in microbial community composition reflect changes in predicted microbial functional pathways. These modules represent a set of manually curated references of metabolic pathways reported to occur in the gut microbiome^52^. Twenty-seven out of 92 (29%) GMMs showed diurnal variation in Chow females, while only 3 out of 92 (3%) GMMs showed significant diurnal rhythms in males (Cosinor analysis, *p* < 0.05) (**Figure 2A, TableS2**).

**Figure 2.**
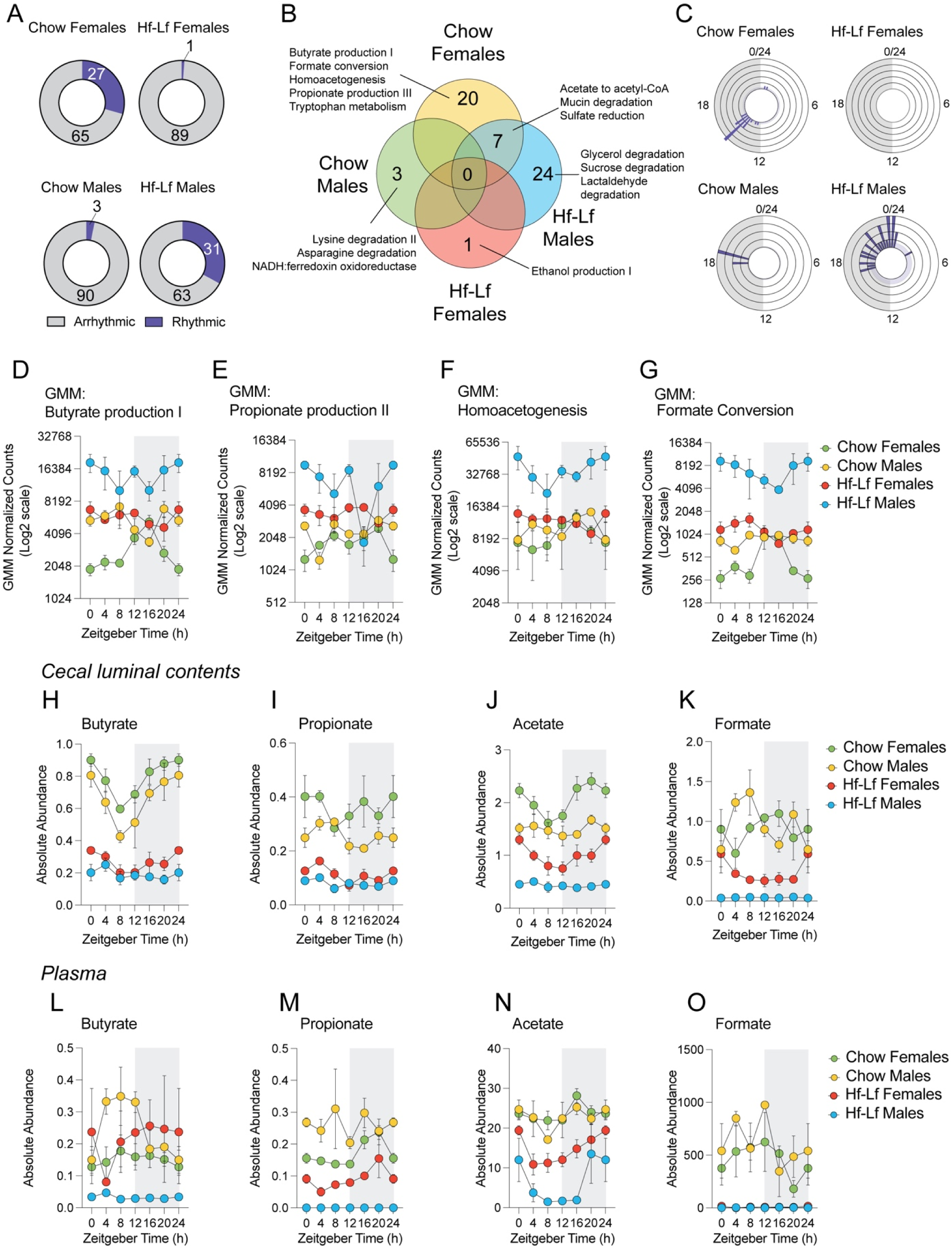
Diet modifies sex differences in diurnal rhythmicity of local and systemic availability of microbial metabolites. A) Oscillation detection of non-rhythmic and rhythmic gut metabolic modules in the cecum of Chow and Hf-Lf mice using cosinor analysis (Cosinor *p* < 0.05). B) Venn diagram depicting differences in rhythmic gut metabolic modules between Chow and Hf-Lf mice. C) Rose plot depicting distribution of maximum availability of rhythmic gut metabolic modules between Chow and Hf-Lf mice. D) Abundance of gut metabolic module: butyrate production I shows sex-specific rhythmicity (Cosinor analysis, Chow females *p* < 0.001; Chow males *p* = 0.257; Hf-Lf females *p =* 0.74; Hf-Lf males *p =* 0.303). E) Abundance of gut metabolic module: propionate production II shows sex-specific rhythmicity (Cosinor analysis, Chow females *p* = 0.021; Chow males *p* = 0.433; Hf-Lf females *p =* 0.98; Hf-Lf males *p =* 0.18). F) Abundance of gut metabolic module: homoacetogenesis shows sex-specific rhythmicity (Cosinor analysis, Chow females *p* <0.01; Chow males *p* = 0.549; Hf-Lf females *p =* 0.44; Hf-Lf males *p =* 0.11). G) Abundance of gut metabolic module: formate conversion shows sex-specific rhythmicity (Cosinor analysis, Chow females *p* <0.001; Chow males *p* = 0.372; Hf-Lf females *p =* 0.17; Hf-Lf males *p =* 0.12). H) Availability of butyrate in cecal lumen (Cosinor analysis, Chow females *p* <0.001; Chow males *p* = 0.372; Hf-Lf females *p =* 0.014; Hf-Lf males *p =* 0.36). I) Availability of propionate in cecal lumen (Cosinor analysis, Chow females *p* = 0.43; Chow males *p* = 0.037; Hf-Lf females *p =* 0.014]3; Hf-Lf males *p =* 0.12). J) Availability of acetate in cecal lumen (Cosinor analysis, Chow females *p* = 0.0006; Chow males *p* = 0.41; Hf-Lf females *p =* 0.006; Hf-Lf males *p =* 0.35). J) Availability of formate in cecal lumen (Cosinor analysis, Chow females *p* = 0.3; Chow males *p* = 0.16; Hf-Lf females *p =* 0.026; Hf-Lf males *p =* 0.79). L) Availability of butyrate in plasma (Cosinor analysis, Chow females *p* = 0.52; Chow males *p* = 0.0027; Hf-Lf females *p =* 0.57; Hf-Lf males *p =* 0.119). M) Availability of propionate in plasma (Cosinor analysis, Chow females *p* = 0.004; Chow males *p* = 0.98; Hf-Lf females *p =* 0.036; Hf-Lf males *p =* 0.117). N) Availability of acetate in plasma (Cosinor analysis, Chow females *p* = 0.35; Chow males *p* = 0.047; Hf Lf females *p =* 0.008; Hf-Lf males *p =* 0.015). O) Availability of formate in plasma (Cosinor analysis, Chow females *p* = 0.077; Chow males *p* = 0.26; Hf Lf females *p =* 0.014; Hf-Lf males *p =* 0.99). Three murine-pathogen free C57Bl/6NTac females and males were used for each condition, for a total of 36 males and 36 females. Acrophases were calculated by cosinor analysis (period = 24 h). Full cosinor analysis in **TableS 2 and 3**. Data is represented as mean ±SEM.

Consistent with the notion that dietary-derived factors entrain diurnal variations of specific taxa, gut metabolic modules that oscillated in Chow mice were abolished in Hf-Lf mice. Cosinor analysis detected that 1 out of 92 (0.01%) GMMs gained diurnal variation in Hf-Lf females, while 31 out of 92 (33%) GMMs gained rhythms in Hf-Lf males (**Figure 2A,C**). Our analysis also revealed sex-specific effects of diet on rhythmic GMMs (**Figure 2B**). For instance, modules that represent synthesis and conversion of primary microbial metabolites, such as formate, butyrate, acetate, and propionate, showed significant oscillations in Chow females but not in males (**Figure 2D-G**). These GMMs remained constitutively high in Hf-Lf females and males regardless of the time of day. Moreover, Chow females and Hf-Lf males showed an overlap in 7 rhythmic GMMs, including processes involved in sulfate reduction and mucin degradation, which is consistent with the bloom in *Desulfovibrio* and Akkermansia observed in Hf-Lf males (see **Figure 1Q,R**). The GMM ethanol production showed diurnal variation in Hf-Lf females, which is consistent with earlier work showing that microbial communities shaped by a Hf-Lf diet exhibit an increased capacity for ethanol production^53^.

### Diet shapes sex differences in the local and systemic availability of microbial-derived metabolites

As our GMM analyses revealed sex and diet-specific differences in the functional potential of microbial pathways involved in the synthesis and conversion of microbial metabolites short-chain fatty acids (SCFAs), we next confirmed whether GMM oscillations reflected diurnal variations in availability of SCFAs in the cecum and plasma^54, 55^. SCFAs are derived from bacterial fermentation of dietary fiber and function as dynamic regulators of host physiology, such as regulating metabolic processes, constraining inflammation, controlling neural circuits involved in feeding and satiety, and regulating gene expression via histone post-translational modifications and inhibition of histone deacetylase activity^56–58^. To measure SCFAs, plasma and luminal cecal contents were collected every 4 hours during a 24-hour period from three female and male mice per time point. We used LC MS/MS to measure the absolute quantification of the SCFAs formate, acetate, propionate, butyrate, valerate, and hexanoate, Valerate and hexanoate fell below our level of detection in this assay. As cecal weight differed across the day in males and females, we normalized cecal SCFA absolute quantities per gram body weight to control for sex differences in cecal SCFA concentration that may be attributed to baseline sexual dimorphism in mouse body weight (**Figure S1**). Plasma SCFA absolute quantities were reported as a concentration (µg/mL) based on volume and thus do not change with body weight^59^.

As predicted by the GMM analysis, diurnal variation in cecal luminal SCFA availability differed in a diet and sex-specific manner (**TableS3**). Chow females showed diurnal variations in the availability of cecal butyrate and acetate, and Chow males in cecal butyrate and propionate availability (Cosinor *p*s < 0.05) (**Figure 2H-J; TableS3**). Formate was available in equal quantities across the day in Chow female and male cecum (**Figure 2K**). Hf-Lf feeding reduced absolute concentration of butyrate, acetate, propionate, and formate relative to Chow mice, suggesting that excess dietary fat, loss of dietary fiber, or a combination of both affects SCFA availability (**Figure 2H-K**). Despite this reduction in cecal SCFA output and amplitude, Hf-Lf mice maintained diurnal variations in SCFA availability in a sex-specific manner. For instance, Hf-Lf females preserved rhythmicity in cecal butyrate and acetate and gained rhythms in cecal propionate, while Hf-Lf males lost rhythmicity in cecal butyrate and propionate (**Figure 2H-K, TableS3**). We replicate earlier work showing loss of diurnal variations in cecal butyrate in male mice^27^, and now show that this previously observed effect is specific to male mice.

We then examined how diurnal variations in cecal luminal SCFA availability influences circulating SCFA levels by analyzing plasma. Propionate was the only SCFA to show diurnal availability in plasma of Chow females, while acetate and butyrate showed rhythmic availability in the plasma of Chow males (**Figure 2L-O; TableS2**). Interestingly, in Chow males, maximum availability of cecal butyrate was ZT21.5 and maximum availability of plasma butyrate was ZT8.5 (**Fig 2. H,L**; **TableS3**) ^60^. Although not significant, a similar trend was observed for propionate, whereby availability of cecal propionate was ZT23.5 and maximum availability of plasma butyrate was ZT4.3 (**Fig 2. I,M**; **TableS3**). Our reconstruction of SCFA suggesting a ∼12h period from cecal derived butyrate to plasma and provide a high-resolution timescale by which specific SCFAs become systemically availability to affect host physiological functions, including regulation of host gene expression, metabolism, and immune function^61^.

Like cecal SCFA availability, Hf-Lf feeding altered diurnal variation in butyrate, propionate, acetate, and formate availability in sex-specific manner (**Figure 2L-O**). Although Hf-Lf reduced absolute abundance in all SCFAs, Hf-Lf females preserved rhythmicity in plasma acetate and propionate, while Hf-Lf males maintained rhythmicity in plasma acetate and butyrate (**Figure 2L, N; TableS3**). As acetate can be synthesized from various diet and microbiota-independent processes^62^, it is thus not surprising that diurnal variations in systemic availability of acetate remains intact in Hf-Lf mice.

Lastly, we examined whether sex differences in diurnal variations of cecal and plasma SCFA availability is dependent on the microbiome. We collected whole ceca and plasma from germ-free female and male mice at ZT6, ZT10, ZT14, and ZT18. Diurnal variations in acetate, butyrate, and propionate availability in the cecum and plasma were abolished and no differences between germ free female and male mice were detected (**FigureS2**). These results confirm that sex differences in diurnal variations in local and systemic SCFA availability are microbiome dependent. Moreover, host genetics and housing conditions also have strong effects on the magnitude of sex differences in mice^63–65^. To determine whether our observed sex differences in diurnal rhythms in microbiota and metabolites may reflect a generalizable pattern in commonly used laboratory mice, we repeated these experiments with BALB/c mice reared in a different animal housing facility. We observed similar sex differences in cecal weight and diurnal rhythms in cecal availability of SCFAs in Chow fed BALB/c female and male mice (**FigureS3**). Overall, our integrated analytical approach identified sex differences in circadian rhythms of microbial structure, composition, predicted metabolic function, and output. The magnitude of sex differences in SCFA availability changed across the day. Furthermore, we show that sex-specific rhythms are determined by various dietary factors, such as the diversity of carbohydrate sources, presence or absence of soluble fibers, and the amount of saturated fats.

### Gut microbiota are required for sex-specific circadian rhythms in host gene expression

Diurnal fluctuations in microbial metabolite availability affect the patterns of gene expression in host tissues^66–68^. This led us to hypothesize that our observed sex differences in the circadian rhythms of gut microbiota and microbial metabolites regulate gene expression of sex-specific cyclic genes within the intestinal tract. We collected small intestinal ileum segments from three female and male Chow-fed conventionalized (CV) and germ-free (GF) mice per time point across a 24-hour period. CV females exhibited diurnal rhythms in 3492 transcripts (∼24%) compared with 278 rhythmic transcripts (2%) in males (p < 0.05 by Cosinor test) (**FigureS4A, TableS4**). Analyses of acrophases revealed sex-specific distribution patterns in the maximum expression of cyclic genes. CV females exhibited a higher number of genes peaking between ZT8 to ZT12 and ZT16 to ZT20, while CV males showed a higher number of genes peaking between ZT2 to ZT6 and ZT12 to ZT18 (**FigureS4B**). Sex differences in the onset and acrophase of feeding have been observed, with females beginning to consume food ∼4hrs earlier than males^69^. This may suggest that phase shifts in gene expression reflect sex differences in the synchronization of feeding rhythms and microbiota, but further testing is needed to confirm these interactions. Additionally, CV females and males showed similar oscillations in only 66 genes, confirming sex differences in intestinal circadian gene expression (**FigureS4C**). The sex difference in the number of rhythmic genes and distribution of peak gene expression was abolished in GF mice (**FigureS4A**). GF females and males showed similar oscillations in 179 genes, indicating that a cluster of genes oscillated in a sex-specific manner in the absence of a microbiome (**FigureS4B**). *Pigv*, a gene that encodes a mannosyltransferase enzyme involved in the biosynthesis of glycosylphosphatidylinositol (GPI), is the only non-sex-specific cyclic gene independent of microbiome status (**FigureS4C**). Defects in GPI biosynthesis can lead to severe morbidity and mortality^70^, highlighting the importance of maintaining rhythmicity independent of sex and microbiome. Thus, GPI serves as a potential candidate gene to further investigate underlying microbiome independent mechanisms that govern circadian rhythms.

We next investigated whether sex-specific rhythmic gene expression is associated with transcriptional networks and functional pathways using Gene Ontology-based overrepresentation analysis (with all pathways having a false discovery rate (FDR) < 0.05). Our analysis revealed sex specific differences in transcriptional pathways involved in metabolism and immunity between CV and GF mice (**FigureS4D; TableS5**). CV female mice exhibited enrichment in pathways involved in intestinal absorption, fatty acid metabolism, cholesterol transport and biosynthesis, gluconeogenesis, immunoglobulin production, and mucosal immune response, while CV males showed enrichment in pathways involved in response to glucose, leptin, and insulin. GF mice did not show enrichment for these biological processes. Specifically, genes enriched in these pathways, including solute carrier family 4 member 1 (*Slac5a1/Slgt1*), free fatty acid receptor 2 (*Gpr42/Ffar2*), CD36 molecule (*Cd36*), leptin (*Lep*), leptin receptor (*Lepr*), insulin receptor substrate 1 (*Irs1*), glucagon-like peptide 1 receptor (*Glp1r*), Ezrin (*Ezr*), and fatty acid binding protein 2 (*Fabp2*) showed cyclicity, and this cyclicity was abolished in GF mice (**FigureS5A-H**). Our results build upon earlier work showing that microbial-derived signals are necessary for generating oscillations in genes involved in intestinal metabolism and immunity^12, 13^. Here, we extend these findings to show that the microbiome is necessary for generating sex-specific circadian rhythms in intestinal gene expression patterns.

### Dietary factors impact circadian rhythms in host gene expression in a sex-specific manner

We next examined whether microbiome-dependent expression of genes involved in metabolism, nutrient uptake, and immunity are influenced by dietary factors^12, 13, 26–28, 71^. We investigated the effect of the Hf-Lf diet on cycling genes in the intestinal tract of female and male mice, comparing them to Chow females and males. Hf-Lf females had fewer rhythmic transcripts (∼28% vs ∼4%), while Hf-Lf males had more (∼7% vs. ∼2%-) compared to the Chow mice (**Figure 3A; TableS6**). The increase in rhythmic genes in Hf-Lf males is likely related to the high degree of inter-sample variability across ZT collection time points in Chow males, which was reduced upon switching to the Hf-Lf diet, as observed in previous studies^49, 50^. Analysis of acrophase distribution shows that the primary loss of rhythmic transcripts occurred during ZT18 to ZT24 in Hf-Lf females compared with Chow females (**Figure 3B**). Hf-Lf males showed a phase advance (∼4 hours) in the acrophase distribution of rhythmic genes, consistent with earlier work on the effects of hypercaloric and energy-dense feeding on circadian rhythms in the liver and adipose tissues^72, 73^ (**Figure 3B**).

**Figure 3.**
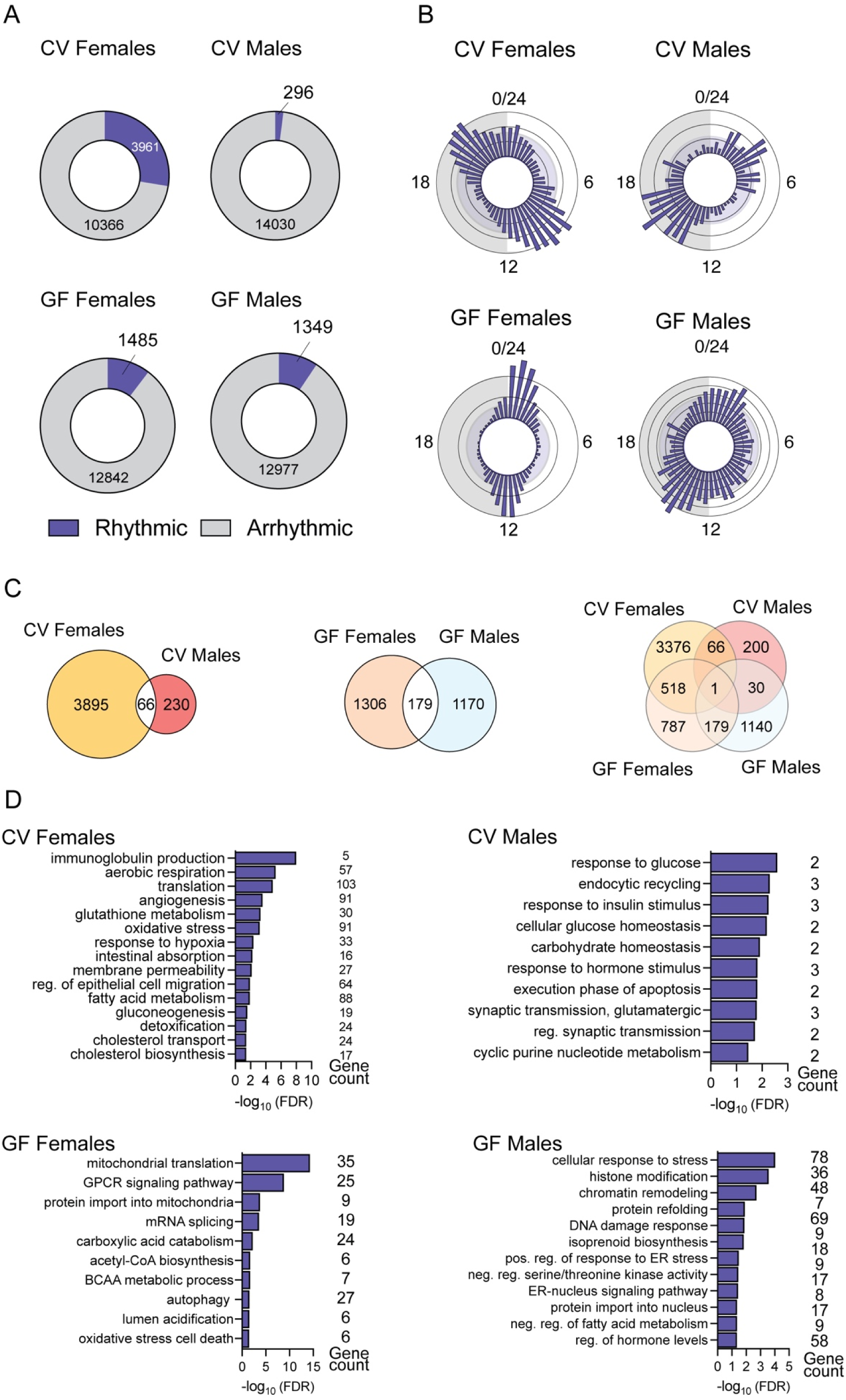
The intestinal microbiota is required for sex differences in diurnal rhythmicity of genes involved in immunity and metabolism. A) Oscillation detection of non-rhythmic and rhythmic transcripts detected in the ileum of Chow diet conventionalized and germ-free female and male mice using cosinor analysis (Cosinor *p* < 0.05) B) Rose plot depicting distribution of maximum expression of rhythmic genes in the ileum of Chow diet conventionalized and germ-free female and male mice C) Venn diagram depicting differences in rhythmic gene expression in Chow diet conventionalized and germ-free female and male mice D) PANTHER overrepresentation test of Gene Ontology using the Biological Process annotation dataset showing enrichment of sex-specific pathways in the ileum of Chow diet conventionalized and germ-free female and male mice (Fisher’s Exact Test, FDR < 0.05). Three murine-pathogen free C57Bl/6NTac females and males were used for each condition, for a total of 30 males and 30 females. Acrophases were calculated by cosinor analysis (period = 24 h). Full cosinor and GO analysis in **TableS 4 and 5**, respectively.

**Figure 4.**
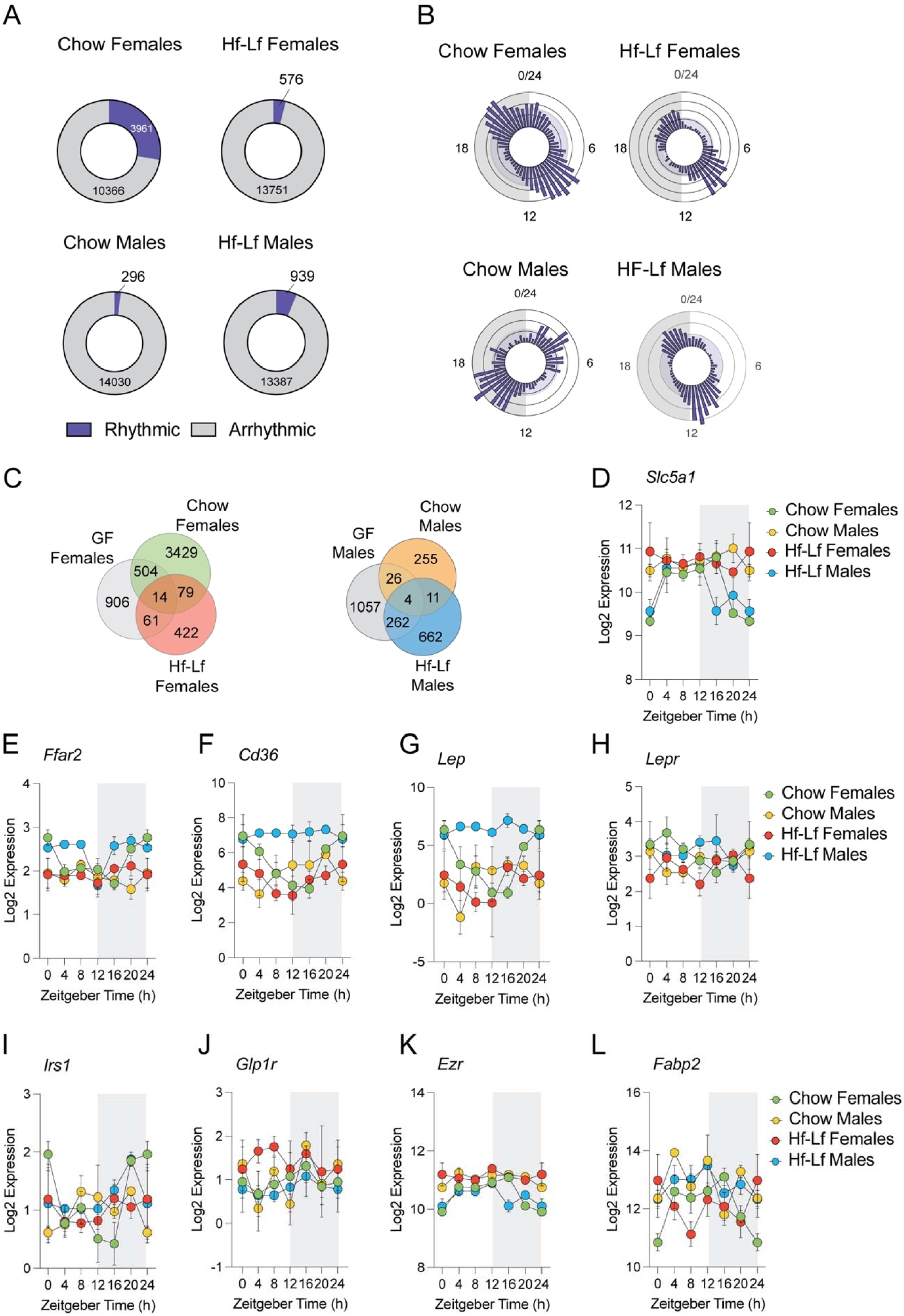
Diet modifies magnitude of sex differences in diurnal rhythmicity of genes involved metabolism. A) Differences in non-rhythmic and rhythmic transcripts detected in the ileum of Chow and Hf-Lf female and male mice using cosinor analysis. B) Rose plot showing differences in acrophase distribution of genes in the adult ileum of of Chow and Hf Lf female and male mice. C) Venn diagram depicting differences in rhythmic gene expression in GF, Chow, and Hf-Lf female and male mice (all *p*s < 0.05 using Cosinor). D) Barplot depicting differences in diurnal variations in sodium-dependent glucose transporter (*Slc5a1)* gene expression in Chow, and Hf-Lf female and male mice (Cosinor analysis, Chow females *p* = 0.014; Chow males *p* = 0.61; Hf-Lf females *p* = 0.93; Hf-Lf males *p =* 0.035). D) Barplot depicting differences in diurnal variations in free fatty-acid receptor 2 (*Ffar2)* gene expression in Chow, and Hf-Lf female and male mice (Cosinor analysis, Chow females *p* = 0.010; Chow males *p* = 0.51; Hf-Lf females *p* = 0.52; Hf-Lf males *p =* 0.15). D) Barplot depicting differences in diurnal variations in the CD36 molecule (*Cd36)* gene expression in Chow, and Hf-Lf female and male mice (Cosinor analysis, Chow females *p* = 0.0009; Chow males *p* = 0.09; Hf-Lf females *p* = 0.21; Hf-Lf males *p =* 0.90). D) Barplot depicting differences in diurnal variations in leptin (*Lep*) gene expression in Chow, and Hf-Lf female and male mice (Cosinor analysis, Chow females *p* = 0.0014; Chow males *p* = 0.13; Hf-Lf females *p* = 0.37; Hf-Lf males *p =* 0.63). D) Barplot depicting differences in diurnal variations in leptin receptor (*Lepr)* gene expression in Chow, and Hf-Lf female and male mice (Cosinor analysis, Chow females *p* = 0.004; Chow males *p* = 0.65; Hf-Lf females *p* = 0.37; Hf-Lf males *p =* 0.90). D) Barplot depicting differences in diurnal variations in insulin receptor substrate 1 (*Irs1)* gene expression in Chow, and Hf-Lf female and male mice (Cosinor analysis, Chow females *p* = 0.011; Chow males *p* = 0.45; Hf-Lf females *p* = 0.72; Hf-Lf males *p =* 0.26). D) Barplot depicting differences in diurnal variations in glucagon-like peptide 1 receptor (*Glp1r*) gene expression in Chow, and Hf-Lf female and male mice (Cosinor analysis, Chow females *p* = 0.63; Chow males *p* = 0.46; Hf-Lf females *p* = 0.79; Hf-Lf males *p =* 0.71). D) Barplot depicting differences in diurnal variations in Ezrin (*Ezr*) gene expression in Chow, and Hf-Lf female and male mice (Cosinor analysis, Chow females *p* = 0.003; Chow males *p* = 0.45; Hf-Lf females *p* = 0.87; Hf-Lf males *p =* 0.15). D) Barplot depicting differences in diurnal variations in fatty-acid binding protein 2 (*Fabp2*) gene expression in Chow, and Hf-Lf female and male mice (Cosinor analysis, Chow females *p* = 0.031; Chow males *p* = 0.65; Hf-Lf females *p* = 0.68; Hf-Lf males *p =* 0.27). Three murine-pathogen free C57Bl/6NTac females and males were used for each condition, for a total of 36 males and 36 females. Acrophases were calculated by cosinor analysis (period = 24 h). Full cosinor analysis in **TableS 6**. Threshold for the cosinor test was set to *p < 0.05*.

Further, a comparison of GF, Chow, and Hf-Lf mice revealed sex-specific effects of diet and the microbiome on oscillating genes (**Figure 3C**). Fourteen genes were shared among all females, and four genes were shared among all males, representing a putative set of sex-specific oscillating genes that are independent of diet and the microbiome. Hf-Lf females gained rhythmicity in 422 genes, and Hf-Lf males gained rhythmicity in 662 genes (**Figure 3C**). As Hf-Lf feeding is known to alter trafficking of metabolites across the intestine, we next examined candidate genes involved in intestinal nutrient absorption and transport that were identified in our comparison between CV and GF mice (**Figure 3E-L**). Hf-Lf feeding resulted in constitutively high expression of *Ffar2*, *Cd36*, and *Lep* while Hf-Lf females-maintained oscillations in these genes, albeit with decreased amplitude (**Figure 3E-F**). *Glp1r* was constitutively expressed in Hf-Lf females relative to Hf-Lf males and Chow mice (**Figure 3J**). The sustained upregulation of genes involved in glycemic control, lipid and fatty acid transport, and leptin signaling may suggest sex-specific metabolic adaptations to Hf-Lf feeding in the intestine^42, 43^. Consistent with this, the Gene Ontology overrepresentation analysis (FDR < 0.05) revealed that Hf-Lf males showed enrichment in gene pathways involved in lipid synthesis and storage, fatty acid metabolism, and response to insulin stimulus. Conversely, Hf-Lf females showed no enrichment in these pathways but a significant enrichment in gene pathways involved in mRNA metabolic processing (**TableS7**).

## DISCUSSION

Sex differences in metabolic functions are crucial for life-long health. Disruption to these processes is associated with sex-specific risk for metabolic, immune, and neural disorders^6^. The gut microbiome is thought to play a key role in meeting the divergent metabolic and immunologic demands of females and males^8, 9^. Further, microbial communities show significant shifts across time-of-day, and disruption to these circadian rhythms is associated with increased risk for metabolic dysfunction and inflammation^14, 22, 30, 31, 34^. Although circadian disruption is often linked to sex-specific health outcomes, studies on circadian rhythms, the microbiome, and health outcomes commonly use only male mice or collapse both sexes into one experimental condition ^35^. To address this knowledge gap, we investigated whether the microbiome is necessary for the maintenance of sex differences in metabolism and transcriptional landscape within the intestinal tract across the time-of-day. Additionally, we examined the sex-specific effect of diet on these host-microbe interactions.

Here we show that the relative abundance of gut microbiota fluctuates across time-of-day in a sex-specific manner. Our analyses focused on the cecal luminal microbiota because of its essential role in producing key microbial substrates important for metabolism ^74^. Consistent with earlier studies examining the fecal microbiota^23^, we observed significant sex differences in the diurnal rhythms in the microbiota diversity, composition, and gut metabolic modules. These sex differences were not uniform across the day, rather, the magnitude of difference between females and males depended on time-of-day. For instance, the relative abundance of Segmented Filamentous Bacteria (SFB) was higher in females at ZT12 (lights off) and males caught up four hours later by ZT16. This rhythmic abundance in SFB drives diurnal rhythms in the expression of antimicrobial peptides as an anticipatory cue for food and exposure of exogenous microbiota during the behaviorally active phase^11^. Considering these observations, our results suggest that sex differences in SFB abundance may reflect sex differences in feeding rhythms. Indeed, a recent report showed that females begin to eat about two hours before lights off, resulting in earlier onset and peak of the feeding rhythm and a phase advancement in overall host-microbe interactions^69^. Given the unique and context-specific ability of SFB to both exacerbate and protect against specific autoimmune related pathologies in mice through alterations to host metabolism and immunity, the possible relationship between sex differences in diurnal variations of SFB, metabolism, immunity, and predisposition to disease warrants further exploration. We also observed sex-specific effects on the microbiome during the behaviorally inactive phase. The relative abundance of *Alistipes* and *Prevotellaceae UCG 001* is increased early in the behaviorally inactive phase in males but not females, while both sexes showed increased abundance of *Lactobacillus*.

The diurnal variation in the relative abundance of cecal luminal microbiota also influenced oscillations of microbial metabolites in the cecum and plasma. We observed a peak in the relative abundance of *Butyricicoccus* which preceded peak availability of butyrate in the cecum, followed by peak availability of plasma butyrate six hours later. While these analyses reconstruct some sex specific oscillations in microbiota and microbial metabolites, the present work remains an underestimate of all oscillating metabolites, given that the microbiome produces tens of thousands of metabolites. Incorporating time-of-day effects on microbiome-metabolite interactions is likely to reveal novel epigenetic, metabolic, and immune associations involved in sex-specific health and disease. Diurnal fluctuations in the microbiome and microbial metabolites are associated with sex specific oscillations in ileal transcriptional networks. Unlike males, females showed stepwise changes in the transcriptional networks across the day, showing time-of-day shifts from pathways involved in defense mechanisms to cholesterol and lipid absorption and epigenetic processes. These distinct shifts in transcriptional networks may reflect the ways in which physiological processes in the intestinal tract are partitioned over the course of the day. Sex differences in host gene expression patterns are abolished in germ-free mice, suggesting that oscillations in microbiota and microbial metabolites are necessary for the distinct transcriptional landscape of the female and male ileum.

Our results touch upon a central question about the evolutionary origins of sex differences in host-microbe interactions. Life history theory posits that evolutionary pressures shape the timing of life events in males and females^75–78^. This theory specifically focuses on life history theory is on defining age-schedules of growth, fertility, senescence, and mortality ^75–78^. As individuals grow and reproduce, they acquire increasing quantities of energy to balance the demands of homeostatic process and reproduction. Thus, the timing of such life events depends on resource and energy substrate availability^75, 77–81^. From the perspective of life history trade-offs, individuals cannot continuously support all biological functions at all times of the day as the energy costs are too high. Circadian rhythms may thus enable individuals to partition and prioritize energy allocation towards life-history events relative to predictable fluctuations in environmental conditions, such as daily fluctuations in food availability ^82^. Recent efforts to integrate the microbiome into life history evolution propose that the composition and function of microbial communities set the pace and timing of life history transitions ^83, 84^. This idea is supported by reports showing that abrupt changes to microbial composition and function coincide with the changes in energy allocations required to transition across development, reproduction, and senescence ^10, 85, 86^. Our work suggests that sex-specific circadian rhythms in host-microbe interactions represent one proximate mechanism that links environmental factors such as food availability to the unique energy demands of female and male life histories.

Another important question concerns the role of diet and the energetic costs needed for the sex-specific maintenance of diurnal fluctuations in microbes, microbial metabolites, and host genes. Earlier work highlighted that the nutritional composition of diets and timing of specific nutrient intake entrain peripheral circadian rhythms in rodents and humans ^87^. For instance, shifting animals from a low-protein, high-carbohydrate diet to a high-protein, low-carbohydrate diet changed rhythmic expression of genes involved in gluconeogenesis in the mouse kidney and liver ^88^. Switching human participants from a high-carbohydrate and low-fat diet to an isoenergetic diet composed of low carbohydrate and high-fat increased the amplitude of rhythmic genes involved in inflammation and metabolism ^89^. One extrapolation of this work is that timing of food intake and the nutritional composition of diets function as entrainment signals for host-microbe interactions. Microbial communities show preferences towards distinct dietary components, including protein, fiber, lactate, and urea^90^. Proportional changes to the accessibility of nutrients result in an ecological advantage for bacteria supplied with their preferred substrate^90^. From this perspective, it is not surprising that transitioning mice from a chow to a high-fat low-fiber diet eliminated circadian rhythms that are entrained by the consumption of a Chow diet. Sex-specific loss of rhythmicity occurred in microbiota and microbial metabolites that utilize soluble fiber, which may suggest that the parallel sex-specific loss of rhythmicity in host genes is related to decreased availability of soluble fiber and warrants further study^54^.

Conversely, the consumption of a high-fat low-fiber diet (Hf-Lf) initiated rhythmic oscillations of microbiota in a sex-specific manner. At the level of the microbiome, Hf-Lf females showed a gain of rhythm in *Blautia* and *Bilophila*, taxa that favors dietary lipids for growth and expansion and associate with visceral fat accumulation^9191, 92^. Hf-Lf males showed gained rhythmicity in the relative abundance of *Bacteroides*, *Akkermansia*, and *Desulfovibrio*. We replicated earlier work showing a gain of rhythm in the hydrogen sulfide-producing taxa *Desulfovibrio*^27^, and we showed that this is a male-specific effect. Female and male mice show distinct metabolic adaptations to Hf-Lf feeding, whereby females, but not males, increase energy expenditure, increase insulin sensitivity, maintain activity levels, and decrease the respiratory quotient, which is suggestive of a greater ability to utilize fat from the diet as an alternative source of energy^42, 43^. Our transcriptomic analysis showing upregulation of genes involved in sensing dietary lipids as well as intestinal lipid uptake and transport in male, but not female, mice may indicate that dysregulation of these genes contributes to weight gain. Indeed, increased intestinal *Cd36, Ffar2,* and *Lep* expression is associated with an increased body mass index^93^. Identification of the underlying mechanisms that maintain rhythmicity of dietary lipid-sensing genes in Hf-Lf females may provide novel targets to reinstate rhythmicity and resistance to diet-induced obesity and metabolic syndrome in males.

In conclusion, our studies provide evidence for sex-specific circadian rhythms in the microbial, metabolite, and transcriptional capacity of the intestinal tract under *ad libitium* feed conditions, indicating sex differences in host-microbe interactions are time-of-day dependent. Our observations also suggest that chronic circadian disruption caused by a high-fat low-fiber diet affects the synchronization between the microbiome, microbial metabolites, and host transcriptome in a sex specific manner. We show that sex-specific risk for diet-induced metabolic dysfunction involves, at least in part, host-microbe circadian rhythms. Furthermore, our findings have practical considerations for studying sex differences in preclinical and clinical settings, as the timing of data collection may influence the detection of sex differences in homeostatic and disease processes^94, 95^. Our reconstruction of sex differences in the onset, acrophase, and amplitude of circadian rhythms can guide the selection of appropriate data collection times. Given the significant effects of the microbiome on health and disease trajectories ^96–101^, further experiments are needed to investigate how disruption to these processes’ influences sex-specific disease risk. Overall, our data highlights the importance of studying sex differences in circadian rhythms across the microbiome, metabolites, and the expression of host genes. A better understanding of these fundamental processes can offer novel insight into diseases that exhibit significant sex-bias in onset, severity, and treatment outcomes.

### Limitations of the study

Gonadal hormones, such as estradiol and testosterone, play a significant role in sex differences, including protection and resistance to diet-induced obesity in female mice^102, 103^. While our results reveal lower variance in females, indicating a limited impact of the estrous cycle on our findings, we did not track the estrus cycle across the 24-hour period collections. Therefore, we cannot exclude the role of gonadal hormone status on interaction between microbial circadian rhythms, diet, host gene expression, and sex differences in metabolism. To obtain additional information on hormone mediated effects on microbial circadian rhythms and host metabolism, we recommend using standard methods to track estrous cycle, manipulate gonadal hormones, and employ the four core genotypes mouse model^69, 102, 103^.

## Acknowledgments

This work is supported by funding from the Eunice Kennedy Shriver National Institute of Child Health and Human Development grants T32HD087194 (PI: SKM) Pilot Project from P50HD096723 (PI: EJ), the National Institute of Diabetes and Digestive and Kidney Diseases grant K01DK1121734 (PI: EJ), a Magee Auxiliary Research Scholars Award (PI: EJ) and Start Up funds from Magee-Womens Research Institute. The Health Sciences Metabolomics and Lipidomics Core is supported by grant S10OD023402 (PI: SGW). We acknowledge Timothy Hand, Jacob DeSchepper, Javonn Musgrove in the University of Pittsburgh Gnotobiotic Animal Studies Facility for technical assistance with circadian collections of germ-free mouse tissues. We acknowledge Will MacDonald in the Children’s Hospital of Pittsburgh Health Sciences Sequencing Core for technical assistance with bulk RNA sequencing on the NextSeq 2000. We acknowledge Heather Evers, Cheyenne Miller, Heather Seiple, Aaron Siegel in the Magee-Womens Research Institute animal facility for technical assistance in care of mice. We acknowledge the University of Pittsburgh Center for Research Computing through the resources provided. Specifically, this work used the HTC cluster, which is supported by NIH award number S10OD028483.

## Author Contributions

Conceptualization, S.K.M., J.P.G., K.E.M., and E.J.; methodology, investigation, and validation, S.K.M., S.J.M., J.P.G., J.K.B., A.L., and A.J.K; formal analysis, S.K.M., A.K., and E.J.; resources, C.A.M., S.G.W., and E.J.; data curation, S.K.M., J.P.G., A.K., and E.J.; writing – original draft, S.K.M., J.P.G., and E.J., writing – reviewing and editing, S.K.M., S.G.W., K.E.M., J.P.G., and E.J.; supervision, E.J.; project administration, J.P.G and E.J.; funding acquisition, E.J.

## Inclusion and Diversity

One or more of the authors of this paper self-identifies as an underrepresented ethnic minority in their field of research or within their geographical location. One or more of the authors of this paper self-identifies as a member of the LGBTQIA+ community. One or more of the authors of this paper received support from a program designed to increase minority representation in their field of research. While citing references scientifically relevant for this work, we also actively worked to promote gender and racial balance in our reference list.

## Materials and Methods

### EXPERIMENTAL MODEL AND SUBJECT DETAILS

#### Animals and tissue collection

All experiments were approved by the University of Pittsburgh and Magee-Womens Research Institute Institutional Animal Care and Use Committee and performed in accordance with National Institutes of Health Animal Care and Use Guidelines. C57Bl/6N female and male mice from Taconic Biosciences (C57Bl/6NTac) arrived at the animal facility aged three weeks. All mice were single housed and acclimated to housing conditions for at least one week before experimentation. All mice were maintained on a 12-hour light/dark cycle (lights on: 0600, Zeitgeber time (ZT) 0; lights off: 1800, ZT 12). Room temperature was maintained between 75-77° F and 40-60% humidity. An Onset HOBO MX2202 Wireless Temperature/Light Data Logger (HOBO Data Loggers, Wilmington, NC) was used to confirm stability of light: dark photoperiod. *Ad libitum* access was provided to water and either a chow diet (NIH #31M Rodent Diet, Envigo; 23.0% protein, 59.0% carbohydrate, 18.0% fat), or a high-fat low-fiber diet (Research Diets D12492; 20.0% protein, 20.0% carbohydrate, 60.0% fat. Germ-free mice were fed chow diet (NIH #31M Rodent Diet, Envigo; 23.0% protein, 59.0% carbohydrate, 18.0% fat). For circadian collections, three mice from each condition from separate cages were euthanized and plasma, ileum, cecum, and brain samples were during a 24h period for each of the 6 timepoints on the Zeitgeber time scale (ZT0, ZT4, ZT8, ZT12, ZT16, ZT20). Due to limitations in after-hours access to gnotobiotic facilities, circadian collections of germ-free mice occurred at 4 timepoints (ZT4, ZT8, ZT12, ZT16). Ileum, cecum, and brain samples were rapidly frozen on dry ice and stored at -80C until further processing.

#### Confirmation of germ-free status

The University of Pittsburgh Gnotobiotic Facility conducts quarterly screening for bacterial contamination in germ-free mice. Screening for the detection of bacterial contamination of mouse feces by aerobic and anaerobic bacteria includes bacterial culture and qPCR assays. Screens were negative on all assays from isolators within which germ-free mouse for this study were housed for both prior to the initiation of experiments and following the conclusion of the experiments. This provides additional confirmation of germ-free status of mice used in these studies.

### METHOD DETAILS

#### Cecal luminal content DNA extraction and 16S rRNA marker gene sequencing

The MagAttract PowerMicrobiome DNA/RNA Kit (Qiagen) extracted genomic DNA from fifty milligrams of cecal luminal contents, using bead-beating on a TissueLyser II (Qiagen), according to the manufacturer’s instructions. 16S libraries were generated using a two-step PCR protocol. Amplicon PCR was performed as follows for amplification of the 16s rRNA V3-V4 region from cecal luminal contents: initial denaturation at 95°C for 3 minutes, following by 25-cyles 95°C for 30 seconds, 55°C for 30 seconds, 72°C for 30 seconds, and a final extension at 72°C for 5 minutes. Resultant 16S V3-V4 amplicons were then purified using AMPure XP beads at a 0.8 ratio of beads to amplicon volume. Illumina Nextera XT Index Primer 1 (N7xx) and Nextera XT Index Primer 2 (S5xx) were used as index primers. Index PCR was performed as follows for amplification of the 16s rRNA V3-V4 region from cecal luminal contents: initial denaturation at 95°C for 3 minutes, following by 8-cyles 95°C for 30 seconds, 55°C for 30 seconds, 72°C for 30 seconds, and a final extension at 72°C for 5 minutes. Results indexed libraries were cleaned up using AMPure XP beads at a 0.8 ratio of beads to indexed library. The concentration of indexed libraries was quantified using Qubit 4 fluorimeter and library fragment size was quantified using an Agilent Tapestation 4200 with D5000 ScreenTapes. Libraries were normalized, pooled, and a paired-end sequencing of pooled libraries was done on an Illumina iSeq 100 System using 2×150bp run geometry in our laboratory.

#### Ileum RNA extraction and preparation for RNA-seq

Frozen tissue samples were homogenized in QIAzol Reagent (Qiagen) using a MiltenyiBiotec gentleMACS Octo Dissociator for 30s. RNA was isolated with Qiagen miRNeasy Mini Kits according to the manufacturer’s instructions. RNA integrity was quantified on an Agilent Tapestation 4200 using TapeStation RNA ScreenTapes. All samples had an RIN score above 8. Sequencing libraries were prepared using Illumina Stranded mRNA prep, Ligation kits with IDT for Illumina RNA UD Indexes Set A, Ligation index adapters. The concentration of indexed libraries was quantified using Qubit and library fragment size was quantified using an Agilent Tapestation 4200 with D5000 ScreenTapes. Sequencing was performed on an Illumina NextSeq 2000 using P3 flow cells and 2×100 paired end-run geometry at the Health Sciences Sequencing Core at Children’s Hospital of Pittsburgh. Sequencing was repeated twice on the same library pool to achieve sufficient resolution and minimize batch effects, producing a yield of an average of 30 – 60 million reads per sample.

#### Quantification of 3NP-Short Chain Fatty Acids

Cecal samples were homogenized with 50% aqueous acetonitrile at a ratio of 1:15 vol: wt. 5µg/mL Deuterated internal standards: (D2)-formate, (D4)-acetate, (D5)-butyrate, (D6)-propionate, (D2) valerate and (D4)-hexanoate (CDN Isotopes, Quebec, Canada) were added. Samples were homogenized using a FastPrep-24 system (MP-Bio), with Matrix D at 60hz for 30 seconds, before being cleared of protein by centrifugation at 16,000xg. Plasma samples were cleared of protein using 4x volumes ice cold 1:1 MeOH: EtOH with vortexing, followed by centrifugation at 16,000xg. 60µL cleared supernatants were collected and derivatized using 3-nitrophenylhydrazine. Each sample was mixed with 20 µL of 200 mM 3-nitrophenylhydrazine in 50% aqueous acetonitrile and 20 µL of 120 mM N-(3-dimethylaminopropyl)-N0-ethylcarbodiimide -6% pyridine solution in 50% aqueous acetonitrile. The mixture reacted at 60_JC for 40 minutes and the reaction was stopped with 0.45 mL of 50% acetonitrile. Derivatized samples were injected (50 µL) via a Thermo Vanquish UHPLC and separated over a reversed phase Phenomenex Kinetex 150mm x 2.1mm 1.7µM particle C18 maintained at 55°C. For the 20-minute LC gradient, the mobile phase consisted of the following: solvent A (water / 0.1% FA) and solvent B (ACN / 0.1% FA). The gradient was the following: 0-2min 15% B, increase to 60%B over 10 minutes, continue increasing to 100%B over 1 minute, hold at 100%B for 3 minutes, reequillibrate at 15%B for 4 minutes. The Thermo IDX tribrid mass spectrometer was operated in both positive ion mode, scanning in ddMS2 mode (2 μscans) from 75 to 1000 m/z at 120,000 resolutions with an AGC target of 2e5 for full scan, 2e4 for ms2 scans using HCD fragmentation at stepped 15,35,50 collision energies. Source ionization setting was 3.0kV spray voltage respectively for positive mode. Source gas parameters were 45 sheath gas, 12 auxiliary gas at 320°C, and 3 sweep gas. Calibration was performed prior to analysis using the PierceTM FlexMix Ion Calibration Solutions (Thermo Fisher Scientific). Integrated peak areas were then extracted manually using Quan Browser (Thermo Fisher Xcalibur ver. 2.7). SCFA are reported as the area ratio of SCFA to the internal standard^104^.

### QUANTIFICATION AND STATISTICAL ANALYSIS

#### Processing and analysis of 16S rRNA marker gene sequencing data

The sequences were demultiplexed on the BaseSpace Sequence Hub using the bcl2fastq2 conversion software (version 2.2.0.) and analyzed using QIIME 2 (version 2022.2)^105^ microbiome bioinformatics platform. Quality control on the resulting demultiplexed forward fastq files were performed using DADA2^106^ denoise-single function with trimming 33bp of the primer sequence. A Naive Bayes feature classifier was trained using SILVA reference sequences^107^ with the q2-feature classifier for taxonomic analysis. The average count per sample was 27,611, with maximum count per sample at 42,407 and minimum count per sample at 10,939. Statistical and meta-analysis of the data was conducted using R phyloseq^108^ and vegan^109^ packages. Data filtering was set to include features where 20% of its values contain a minimum of four counts. In addition, features that exhibit low variance across treatment conditions are unlikely to be associated with treatment conditions, and therefore variance was measured by interquartile range and removed at 10%. Data were normalized by using trimmed mean of M-values. Taxa identified as cyanobacteria or ‘unclassified’ to the phylum level were removed. Oscillation of microbiota abundance and period of oscillation were detected by cosinor analysis using the R package DiscoRhythm^110^. Taxa with p < 0.05 over a 24-h oscillation period are reported based on Cosinor analysis.

#### Processing and analysis of bulk RNA-seq data

Concatenated FASTQ files generated from Illumina were used as input to kallisto^111^, a program that pseudoaligns high-throughput sequencing reads to the *Mus musculus* reference transcriptome (version 38) and quantifies transcript expression. We used 60 bootstrap samples to ensure accurate transcript quantification. Gene isoforms were collapsed to gene symbols using the Bioconductor package tximport (version 3.4). Genes were filtered to counts per million >1 in at least three samples. The filtered gene list was normalized using trimmed mean of M-values in edgeR^112^. Oscillation of microbiota abundance and period of oscillation were detected using Cosinor analysis using the R package DiscoRhythm^110^. Transcripts with p < 0.05 over a 24-h oscillation period are reported. Over-representation analysis of Gene Ontology: Biological Process terms to identify enriched molecular pathways/processes in both the top enriched and top rhythmic gene lists was performed with clusterProfiler^113^ and PantherDB^114^.

#### Analysis of targeted metabolomics data

Oscillation of absolute or relative metabolite abundance and period of oscillation were detected using Cosinor analysis using the R package DiscoRhythm^110^. Metabolites with p < 0.05 over a 24-h oscillation period are reported.

#### Quantification and statistical analysis

Statistical information including sample size, mean, and statistical significance values are shown in the text or the figure legends. A variety of statistical analyses were applied, each one specifically appropriate for the data and hypothesis, using the R statistical environment. For standard metabolic endpoints, analysis of variance (ANOVA) testing with repeated-measures corrections and Bonferroni post-hoc tests were used, with significance at an adjusted plJ<lJ0.05. Processing of RNA-Seq data was conducted using standardized and published protocols. GraphPad Prism and Adobe Illustrator were used for generating figures. No custom script was used to analyze RNA sequencing data.

#### Statistical analysis of data

Number of samples used per time point and condition are described under ‘Animals and tissue collection’ section and in the caption of each associated figure along with the statistical method used for analysis. All analyses were performed in python version 3.6.12 (Python Software, 2020) or R version 4.1.0 (R Core Team, 2021)

### STAR METHODS

**Table 1.**
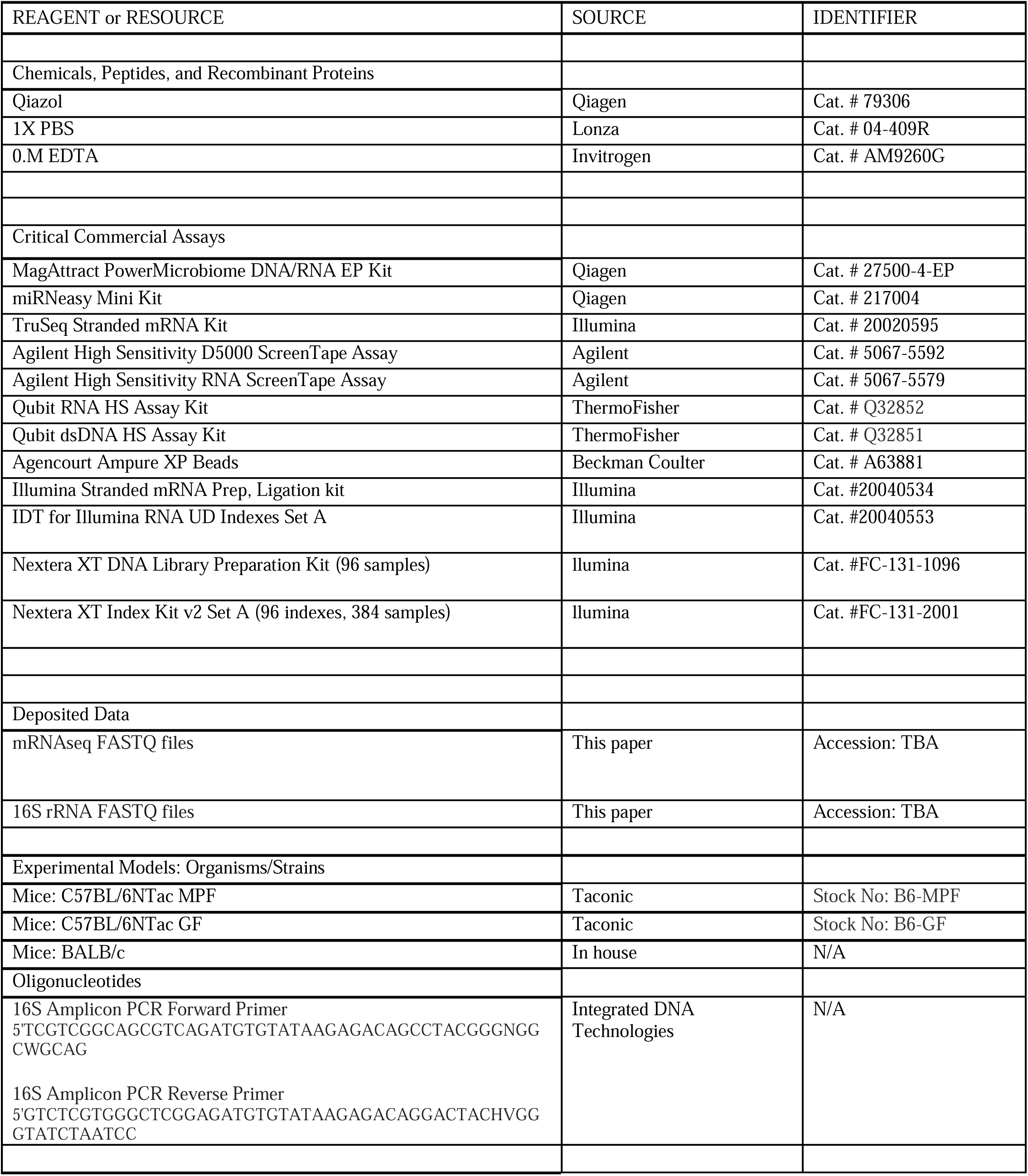
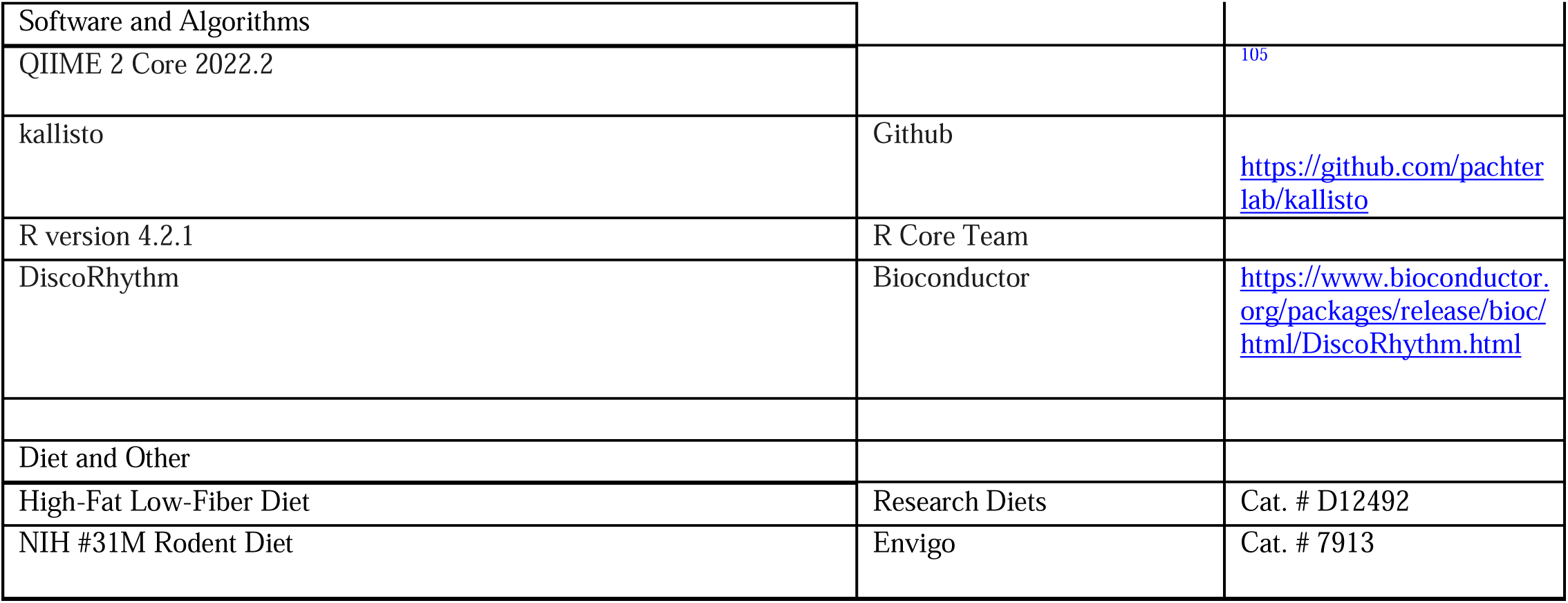
Reagents and Resources used in this report.

### SUPPLEMENTALFIGURES

**Supplemental Figure 1.**
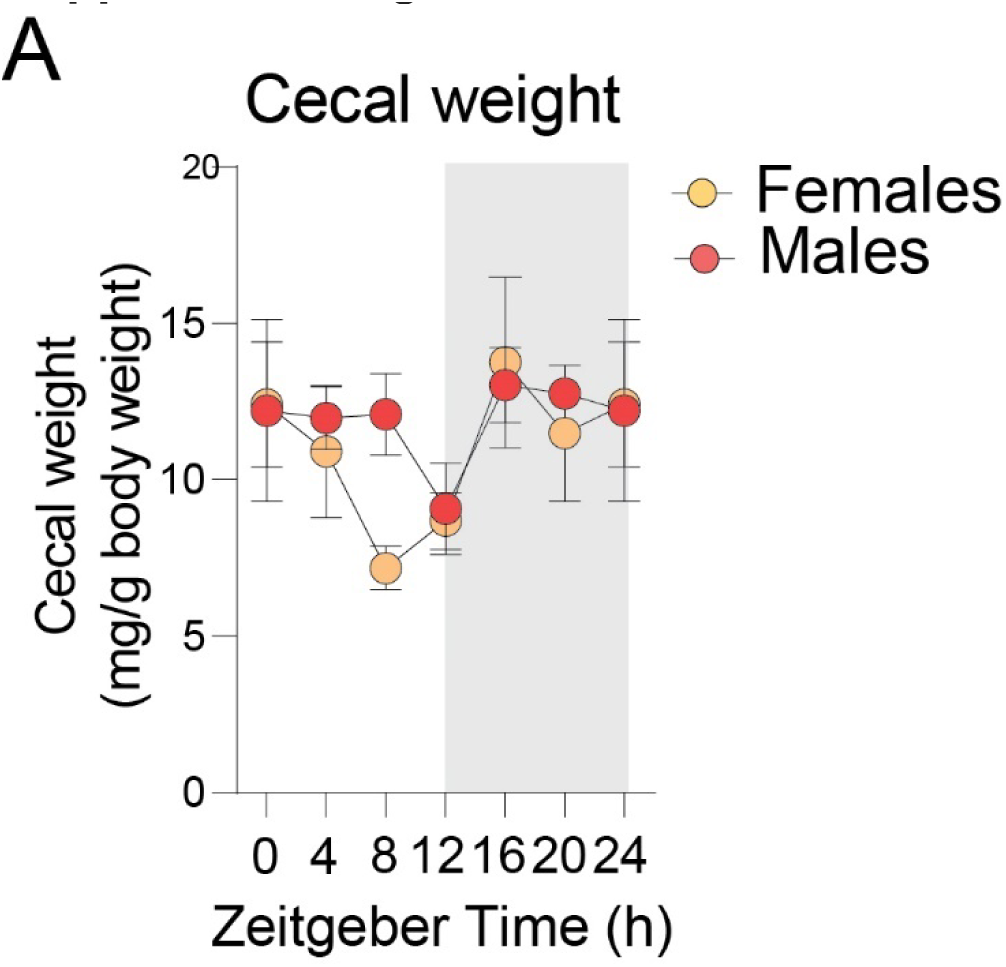
Cecal weight differs by time-of-day and by sex. A) Cecal weights relative to total body weight in Chow and Hf-Lf mice across timepoints. Three murine-pathogen free C57Bl/6NTac females and males were used for each condition, for a total of 18 males and 18 females. Data is represented as mean ± SEM.

**Supplemental Figure 2.**
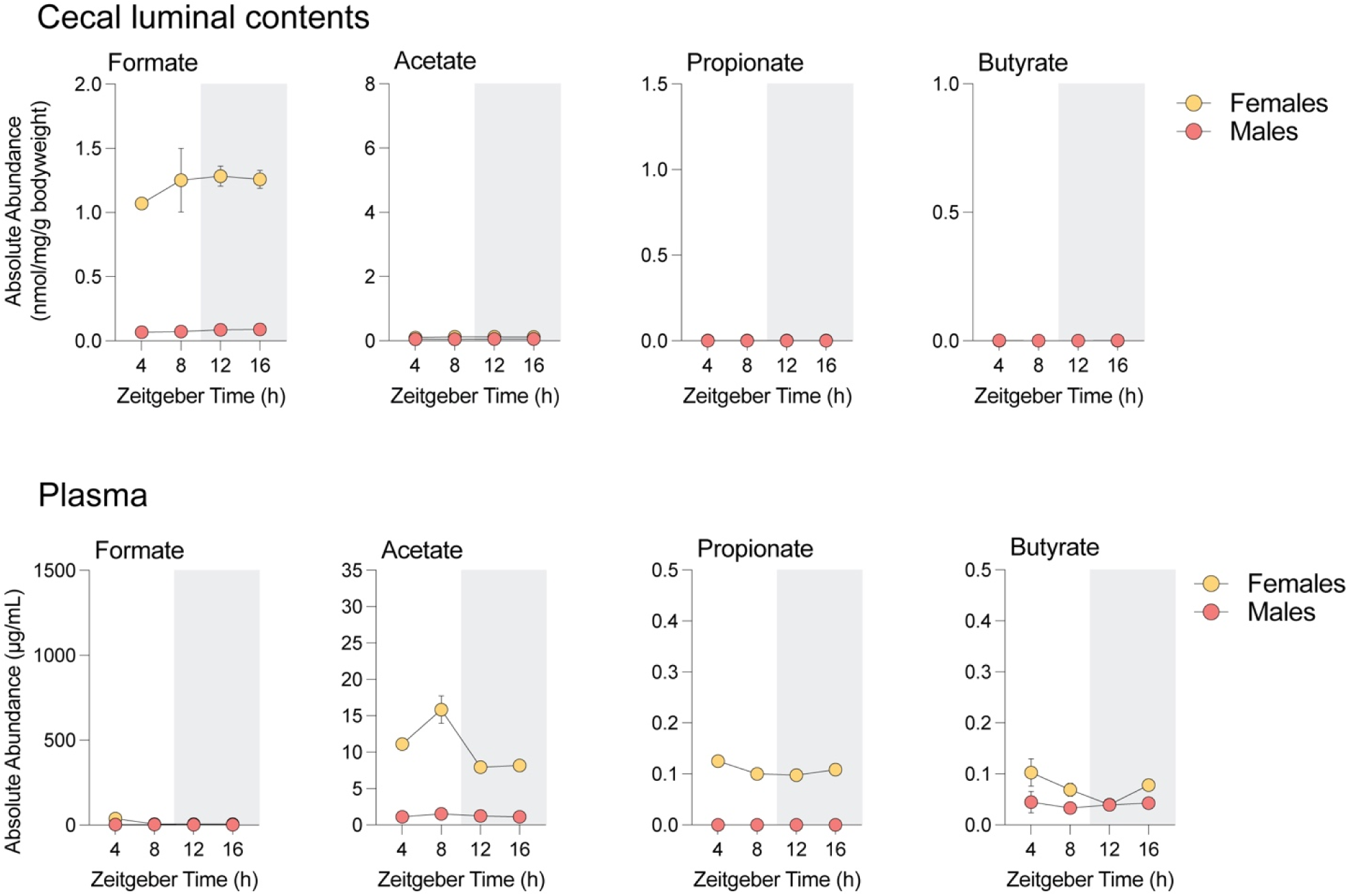
Gut microbiota is necessary to maintain diurnal variations in microbial metabolite availability in germ-free mice. A) Availability of formate, acetate, propionate, and butyrate in cecal lumen shows loss of sex-specific rhythmicity in Germ-Free mice (all *p*s > 0.05 using Cosinor analysis) B) Availability of formate, acetate, propionate, and butyrate in plasma shows loss of sex-specific rhythmicity in Germ-Free mice (all *p*s > 0.05 using Cosinor analysis). Three Germ-free C57Bl/6NTac females and males were used for each condition, for a total of 12 males and 12 females. Data is represented as mean ± SEM.

**Supplemental Figure 3.**
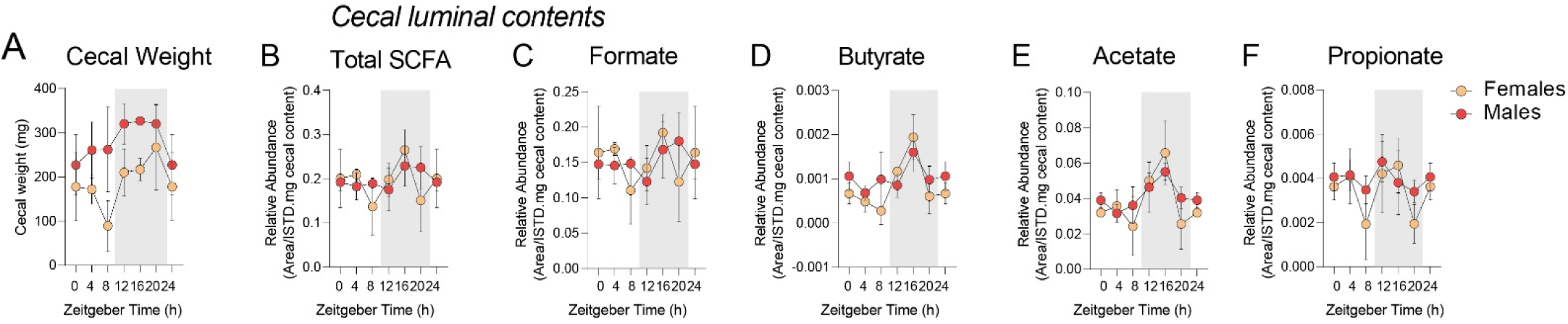
Sex differences in rhythmic availability IN BALB/c mice. A) Cecal weights in female and male BALB/c mice across timepoints. B-F) Quantification of short-chain fatty acids in female and male BALB/c mice across a 24-hr period, with samples collected every 4hrs. Three BALB/c females and males were used for each condition, for a total of 18 males and 18 females. Data is represented as mean ± SEM.

**Supplemental Figure 4.**
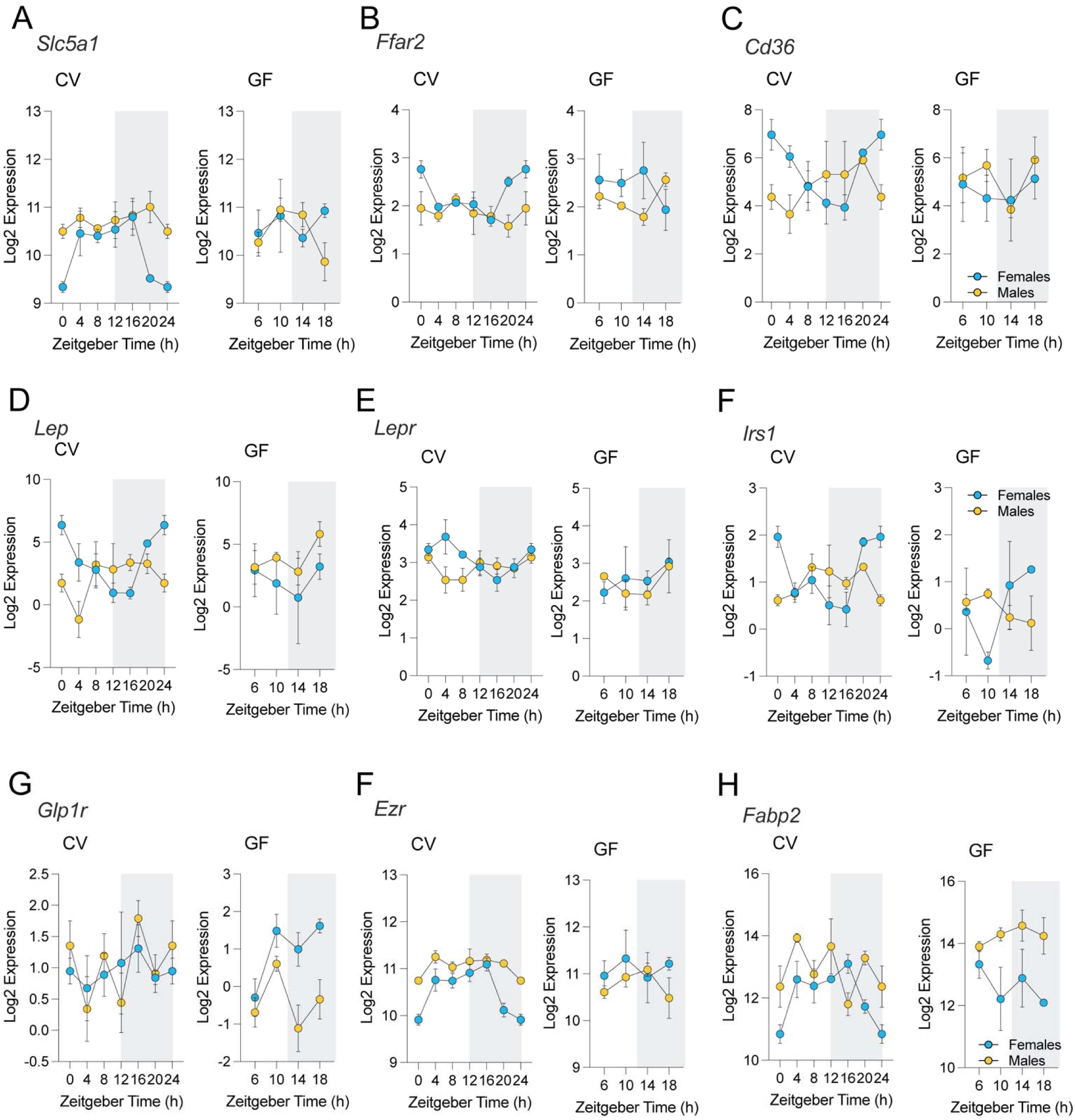
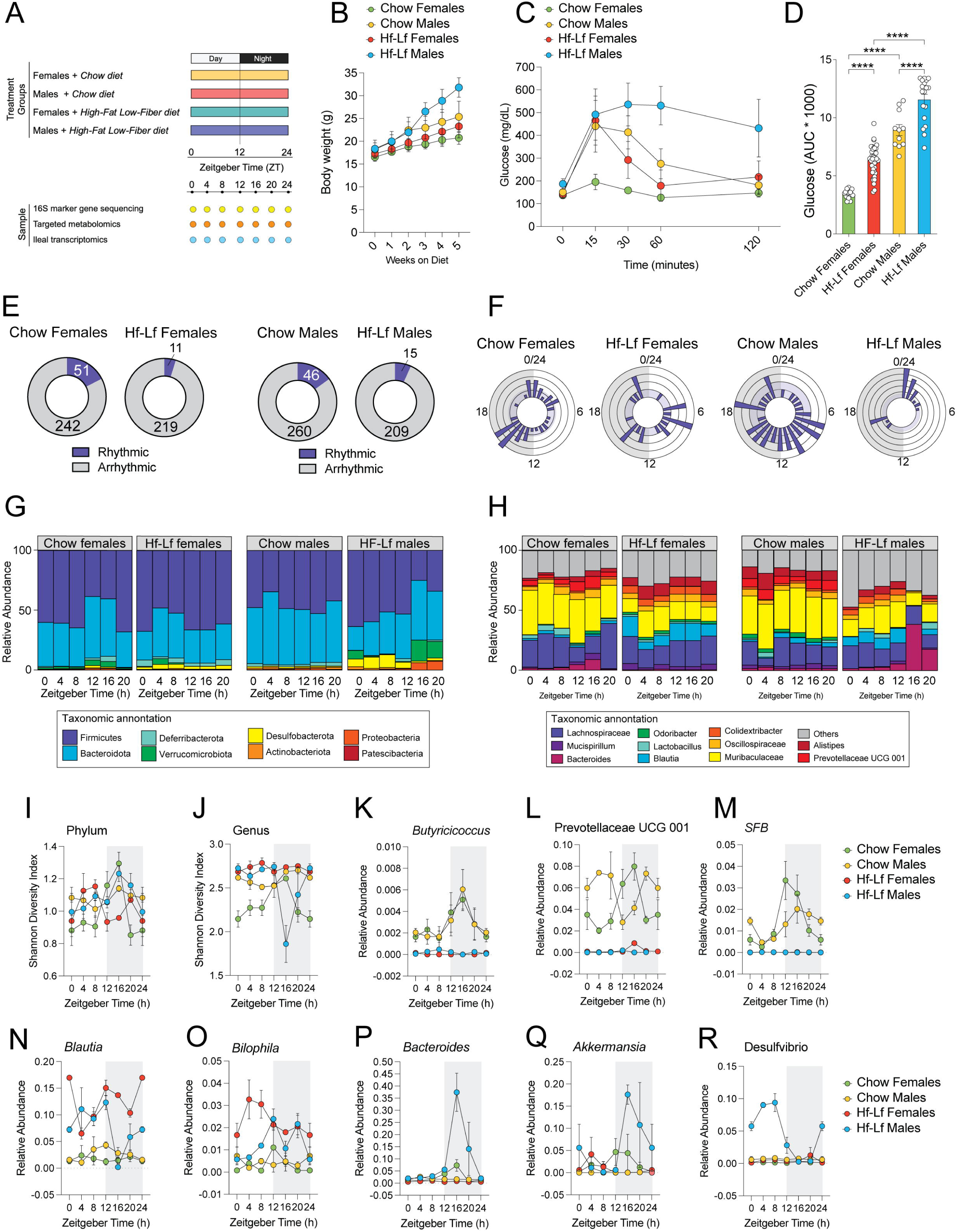
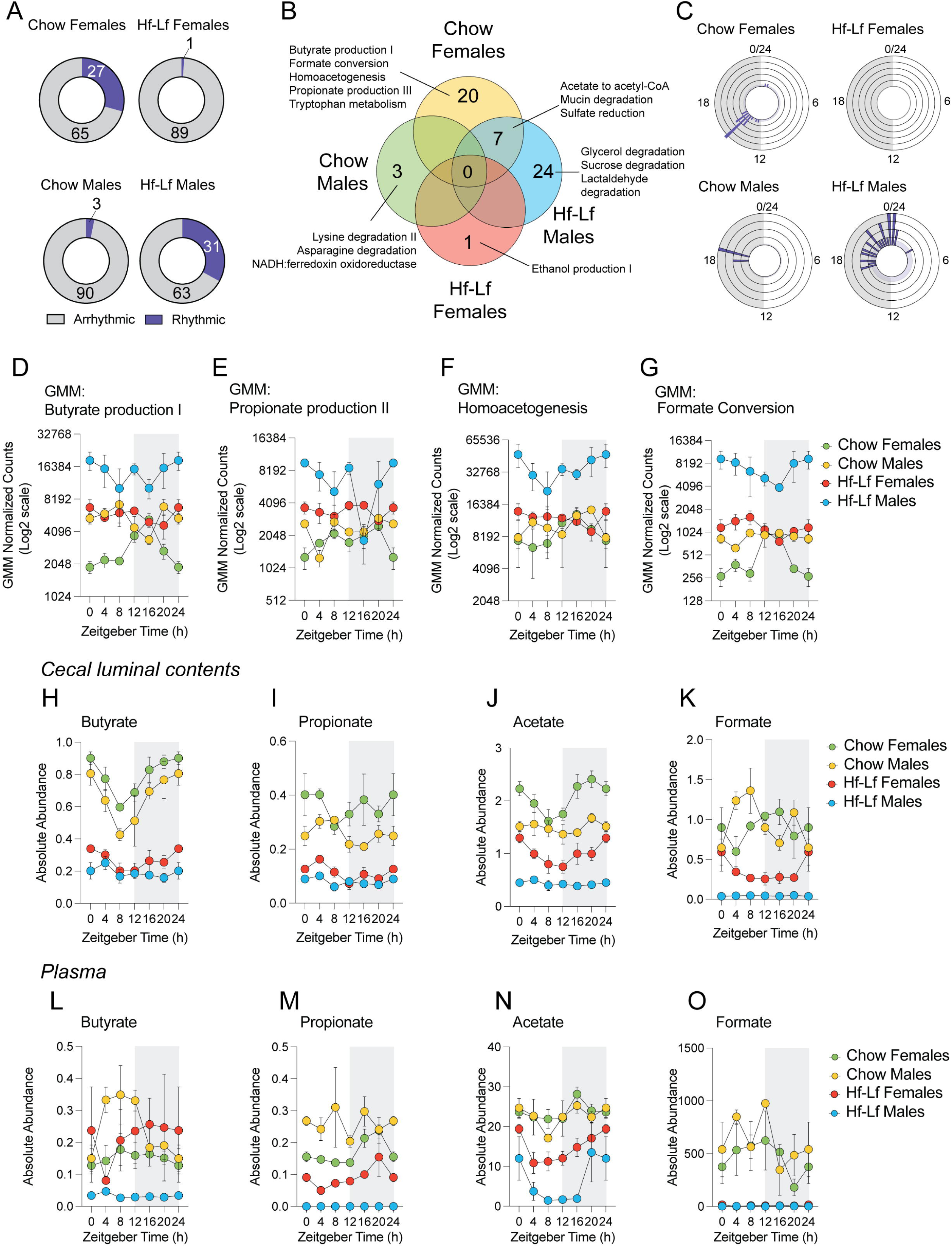
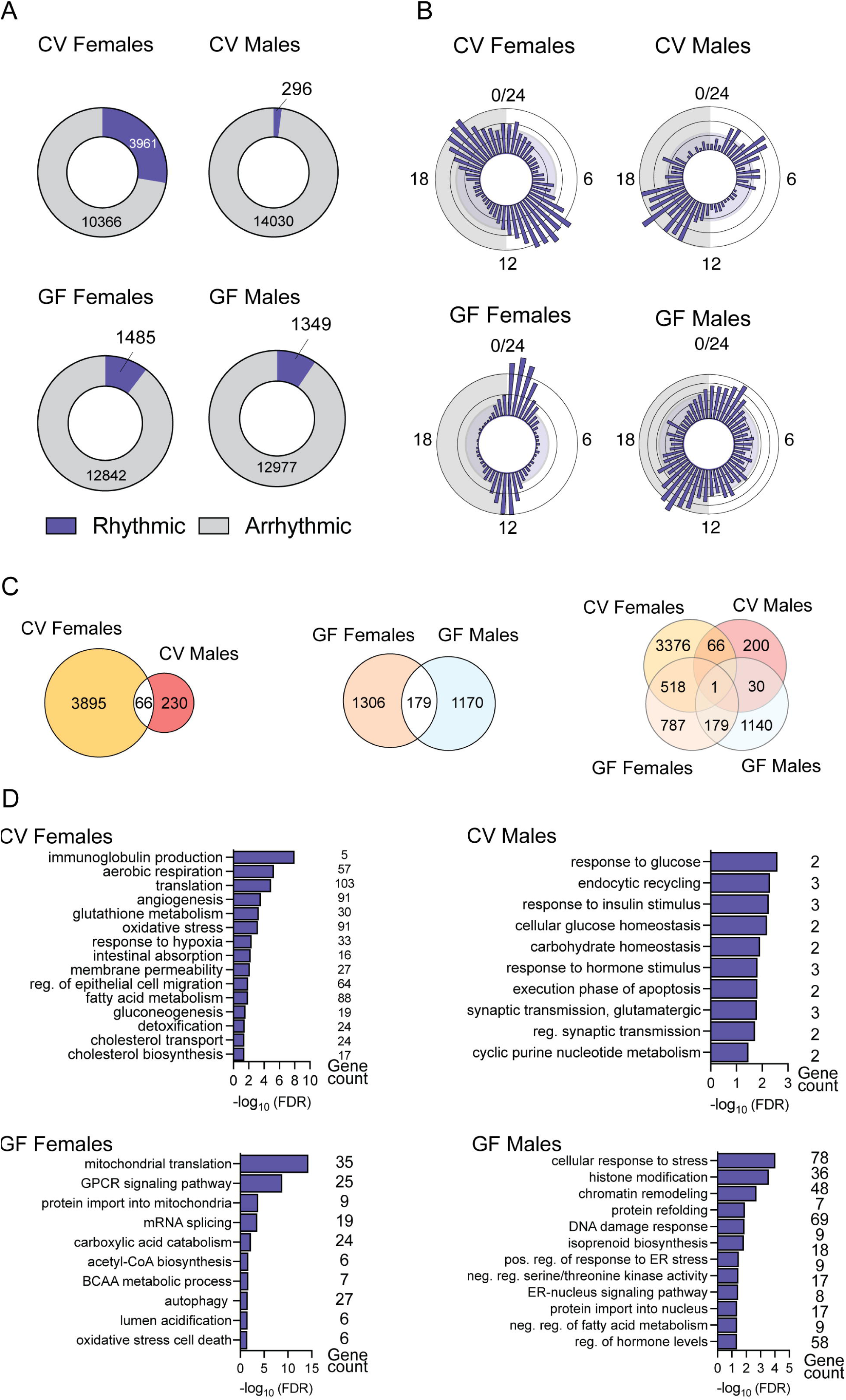
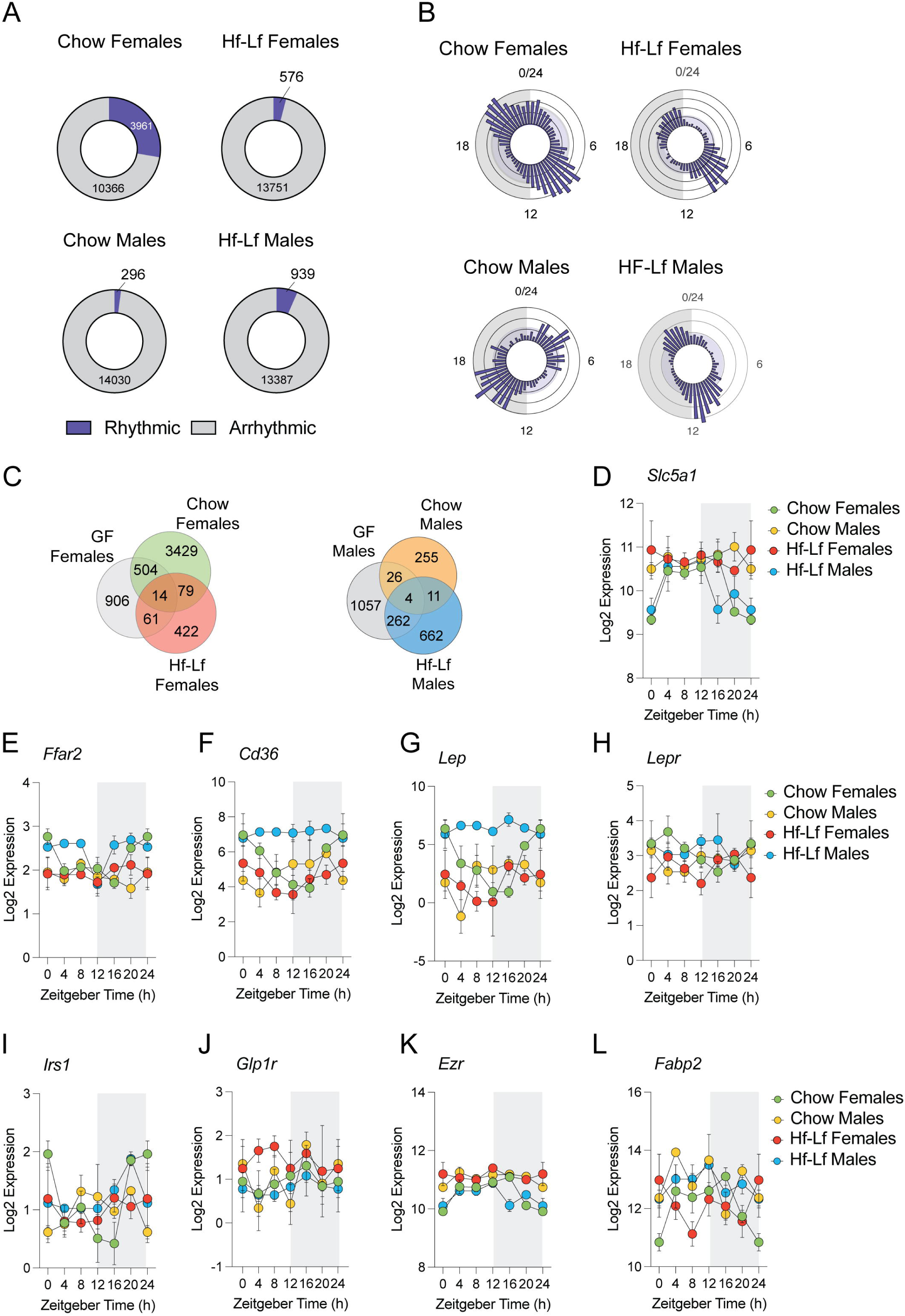
Gut microbiota is necessary to maintain diurnal variations in intestinal genes involved in metabolism and nutrient absorption. A) Barplot depicting differences in diurnal variations in sodium-dependent glucose transporter (*Slc5a1)* gene expression in CV and GF female and male mice (Cosinor analysis, CV females *p* = 0.014; CV males *p* = 0.61; GF females *p* = 0.704; GF males *p =* 0.38). B) Barplot depicting differences in diurnal variations in free fatty-acid receptor 2 (*Ffar2)* gene expression in CV and GF female and male mice (Cosinor analysis, CV females *p* = 0.010; CV males *p* = 0.51; GF females *p* = 0.21; GF males *p =* 0.48). C) Barplot depicting differences in diurnal variations in CD36 molecule (*Cd36)* gene expression in CV and GF female and male mice (Cosinor analysis, CV females *p* = 0.0009; CV males *p* = 0.09; GF females *p* = 0.42; GF males *p =* 0.57). D) Barplot depicting differences in diurnal variations in leptin (*Lep*) gene expression in CV and GF female and male mice (Cosinor analysis, CV females *p* = 0.0014; CV males *p* = 0.13; GF females *p* = 0.46; GF males *p =* 0.086). E) Barplot depicting differences in diurnal variations in leptin receptor (*Lepr)* gene expression in CV and GF female and male mice (Cosinor analysis, CV females *p* = 0.004; CV males *p* = 0.65; GF females *p* = 0.74; GF males *p =* 0.36). F) Barplot depicting differences in diurnal variations in insulin receptor substrate 1 (*Irs1)* gene expression in CV and GF female and male mice (Cosinor analysis, CV females *p* = 0.011; CV males *p* = 0.45; GF females ; GF males *p =* 0.48). G) Barplot depicting differences in diurnal variations in glucagon-like peptide 1 receptor (*Glp1r*) expression in CV and GF female and male mice (Cosinor analysis, CV females *p* = 0.63; CV males *p* = 0.46; GF females *p* = 0.35; GF males *p =* 0.72). H) Barplot depicting differences in diurnal variations in Ezrin (*Ezr*) gene expression in CV and GF female and male mice (Cosinor analysis, CV females *p* = 0.003; CV males *p* = 0.45; GF females *p* = 0.68; GF males *p* = 0.27). I) Barplot depicting differences in diurnal variations in fatty-acid binding protein 2 (*Fabp2*) gene expression in CV and GF female and male mice (Cosinor analysis, CV females *p* = 0.031; CV males *p* = 0.65; GF females *p* = 0.71; GF males *p =* 0.31). Three Germ-free C57Bl/6NTac females and males were used for each condition, for a total of 12 males and 12 females. Full cosinor analysis in TableS 4. Data is represented as mean ± SEM.

